# Membrane sialylation orchestrates cellular gateways: A spatiotemporal analysis of cellular transport using DNA nanocages via membrane charge modulation

**DOI:** 10.64898/2026.05.05.722926

**Authors:** Geethu Prakash, Bhagyesh Parmar, Harsh Dave, Sivaraman Dhanasekaran, Dhiraj Bhatia

## Abstract

Negatively charged DNA nanostructures, such as tetrahedral nanocages, are internalized by cells despite the electrostatic repulsion from the anionic cell membrane, and, paradoxically, cancer cells, which carry intrinsically higher negative charge due to overexpression of sialic acids on their cell surface, show markedly higher uptake than normal cells. This contradiction exposes a fundamental gap in our understanding of how these anionic nanostructures overcome this repulsion. Using chemical modulation of cell-surface sialylation in RPE1 cells to create three groups with altered sialylation levels, together with inhibitor-based dissection of endocytic pathways, we demonstrate that an increase in cell surface sialylation governs the uptake of DNA tetrahedra not through electrostatics but by structurally remodeling the cell membrane via rearrangement of the GM1 lipid raft microdomain, recruiting caveolae-mediated endocytosis as an additional pathway alongside clathrin-mediated endocytosis, thereby increasing the intake of the nanostructure. These findings reframe tumor hyper-sialylation as a determinant of the uptake of anionic nanostructures, such as DNA tetrahedra, and as a targetable parameter for rational optimization of DNA-based nanotherapeutics against cancer.

**Graphical abstract:** 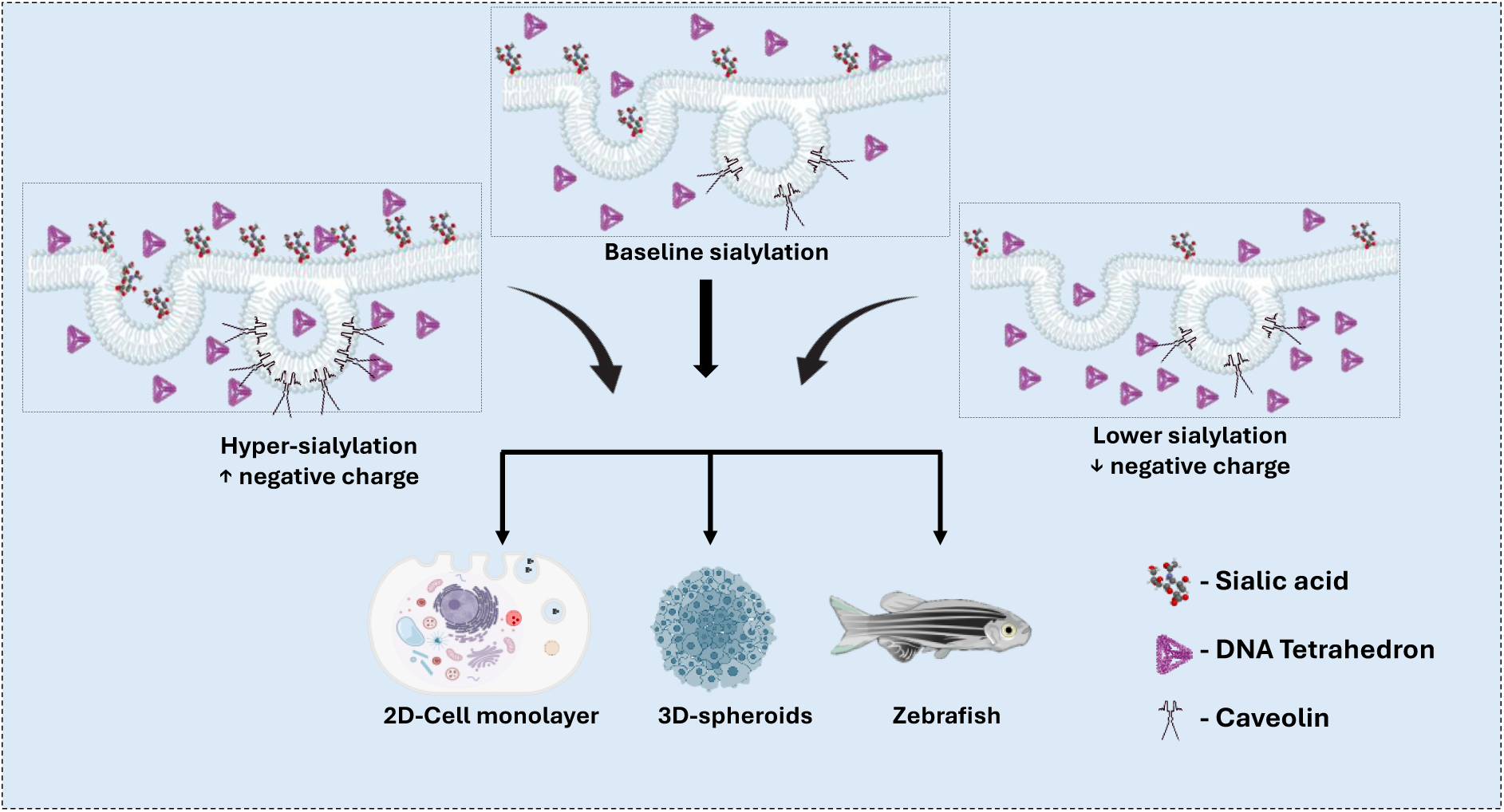

## Main

The cell membrane is a dynamic, highly regulated interface that selectively controls the entry of molecular and nanoscale entities into the cell. For any therapeutic or diagnostic cargo to enter the cell and exert its intracellular function, successful engagement with this barrier is the first and most critical determinant. To date, nanoparticles have revolutionized this challenge by enabling tunable size, shape, and surface chemistry to enhance cellular uptake for applications ranging from drug delivery to biosensing and imaging. Among these, DNA tetrahedral nanocages (TDNs) are particularly attractive because of their compact, rigid architecture, well-defined geometry, and other features, which together support efficient cellular internalization across diverse cell types compared to other DNA nanocages [1].

At physiological pH, DNA possesses a dense negative charge due to its phosphate backbone, while the mammalian cell surface is similarly anionic. This intrinsic negative charge on the mammalian cell surface arises from the glycolipids and glycoproteins [2]. The negative charge on these entities is, in turn, primarily contributed by a nine-carbon monosaccharide called sialic acid, which terminates the glycan chains on cell-surface glycoproteins and glycolipids [3]. Beyond this, sialic acid residues mediate critical processes, including cell-cell recognition, signaling, and immune regulation[4]. In pathological contexts, particularly in cancers, aberrant over-expression of sialic acids, termed as hyper-sialylation, is a hallmark that drives tumor progression, metastasis, and immune evasion [5], [6].

The dense negative charge on both DNA and the cell membrane naturally predicts strong mutual repulsion that should block membrane association and uptake. Paradoxically, DNA TDNs are efficiently internalized across diverse cell types, with surprisingly enhanced TDN uptake in hypersialylated cancer cells compared to non-cancerous cells with baseline sialylation. This discrepancy exposes a fundamental gap in understanding these interactions and raises a central question: why do cancer cells exhibit enhanced uptake of negatively charged DNA TDNs despite their highly negative, sialic acid-rich surface? Here, we hypothesize that pathological hyper-sialylation does not simply create a more repulsive barrier but instead establishes a permissive membrane landscape that actively favors the internalization of DNA nanocages. To understand this, we constructed RPE1 cell groups with three different sialylation states: a baseline, untreated control group; a hypersialylated group mimicking the sialylation levels of cancer cells; and a desialylated group with comparatively lower sialylation levels. This model serves as a baseline reference to provide a physiologically relevant control for studying the effects that truly arise from hyper-sialylation, since direct comparison with cancer cells is confounded by multiple oncogenic alterations that affect not only sialylation but also receptor expression, etc. Systematic quantification of TDN uptake across this model revealed that the electrostatic barrier is not the dominating determinant of TDN uptake. Instead, hyper sialylation expands GM1-rich lipid raft microdomains at the cell surface, making caveolae-mediated endocytosis an additional uptake pathway for TDN, along with clathrin-mediated endocytosis. Validation of this trend in 3D spheroid and zebrafish models confirmed that the glycocalyx signature that cancer cells acquire for immune evasion rewires endocytic mechanisms, thereby enhancing DNA TDN uptake. Cell surface sialylation is not, therefore, a mere electrostatic feature of cells, but an active determinant of DNA nanocage engagement and entry into cells.

### DNA tetrahedra form monodisperse, structurally defined nanocages for uptake studies

A precise examination of the determinants of nanocage uptake requires a nanostructure population that is compositionally and dimensionally uniform. Among the different DNA nanostructures constructed so far, DNA TDNs are more compact, rigid, with well-defined geometry, and efficient cellular uptake across different cell types. We therefore utilized DNA TDN structures for our study. The construction of DNA TDNs utilized a step-down thermal annealing strategy using four single-stranded complementary oligonucleotide strands. Equimolar concentrations (10µM) of the four oligonucleotides were subjected to an initial denaturation at 95°C for 30 minutes, followed by a 5°C reduction at each step for 15 minutes till 4°C, in the presence of 2mM MgCl_2_ (**Figure 1.a**). Successful assembly of the TDN was confirmed by an Electrophoretic Mobility Shift Assay (EMSA) on a 10% polyacrylamide gel. EMSA revealed progressive mobility retardation with an increase in the number of participating oligonucleotides and confirmed assembly of DNA TDN (**Figure 1.b**). Dynamic light scattering (DLS) measurement in nuclease-free water at 25°C indicated an average hydrodynamic diameter of 10nm (**Figure 1.c**). Zeta potential of TDN in nuclease-free water was measured at -18.84 mV, indicating the anionic nature of TDN contributed by the phosphate backbone on the oligonucleotides (**Figure 1.d**). Atomic force microscopy (AFM) imaging further confirmed the formation of DNA TDN at higher resolution (**Figure 1.e**). To enable fluorescent quantification of TDN within cells, one of the four oligonucleotides in the nanostructure was labeled with Cy3 or Cy5. The stability of DNA TDNs is crucial in cellular environments, since degradation of the nanostructure can give deceptive fluorescent signals. Therefore, the stability of TDNs was studied in serum-free media and 10% Fetal Bovine Serum (FBS) at 37°C and in E3 media (for Zebrafish studies) at 28°C for up to 24 hours. Structural integrity was defined as ≥80% retention of the initial intensity. TDN maintained integrity for 1 hour in serum-free media, FBS, and E3 (**Figure S.1**). The assay validates the use of TDN in physiological conditions over the timescale employed in uptake experiments.

**Figure 1:**
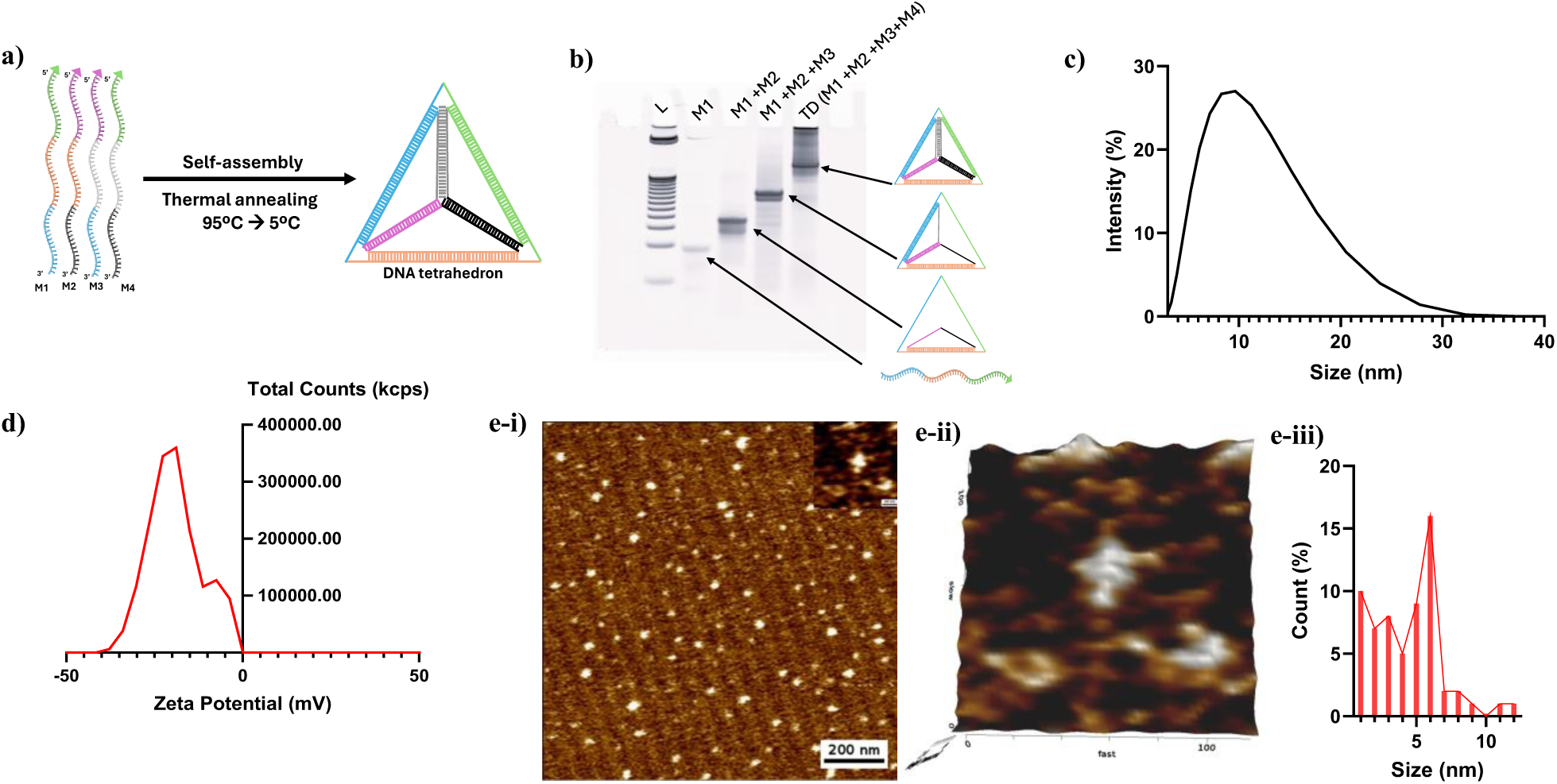
Synthesis and characterization of DNA TDN. a) Schematic illustration of TDN assembly from four single-stranded oligonucleotides (M1 to M4) via thermal annealing. b) EMSA showing the retardation in the mobility of resulting 3D structures with an increase in the number of participating oligonucleotides. Lane 1, DNA ladder (25bp); lane 2, single ssDNA strand (M1); lane 3, formation of a duplex structure with two ssDNA strands (M1 and M2); lane 4, participation of three ssDNA strands (M1, M2, and M3), and lane 5 indicates the TDN structure with four ssDNA strands (M1, M2, M3, and M4). c) Analysis of the hydrodynamic diameter of TD using Dynamic Light Scattering (DLS) indicates that the hydrodynamic diameter of TDN is ∼10nm, and d) the zeta potential is -18.84mV. e) AFM images showing e-i) the topology of TDN (2D), e-ii) 3D AFM image of TDN and e-iii) size distribution profile of TDN measured using AFM.

### Chemical glycoengineering generates three distinct membrane sialylation states without inducing cytotoxicity

To understand the contribution of cell membrane sialylation to DNA TDN uptake, we constructed three distinct groups of Retinal Pigment Epithelial (RPE1) cells with different levels of membrane sialylation: (i) Control cells maintained in standard Dulbecco’s Modified Eagle’s Medium (DMEM) with 10% FBS, representing the physiological baseline sialylation of normal epithelial cells; (ii) hypersialylated cells generated by supplementing DMEM media with 2mM of N-acetylneuraminic acid (Neu5Ac), a sialic acid donor, for 96 hours, exploiting the capacity of cells to incorporate exogenously supplied sialic acids into glycoproteins and glycolipid terminals via the salvage biosynthetic pathway [7]. This mimics the hypersialylated states of cancer cells and (iii) desialylated cells generated by supplementation of 3Fax-Peracetyl Neu5Ac (a sialyltransferase inhibitor; 2µM) in DMEM media to overcome intrinsic sialylation followed by incubation with α-2-3,6,8-neuraminidase from Clostridium perfringens (0.1U/ml) for 30 minutes to cleave intrinsic sialic acid on cells [8], [9]. This confirmed near-complete removal of sialic acid from cells without causing cytotoxicity. Final concentrations of each treatment were decided after viability assessment and optimization protocols (**Figure S2, S3, and S4**).

Surface sialylation of all three groups was quantified using wheat germ agglutinin (WGA), a lectin-specific, cell-impermeant dye that labels Neu5Ac residues on the cell surface [10], [11]. Alexa 647 labeled WGA staining followed by confocal microscopy revealed alterations in sialic acid content on the cell surface of the three groups. Mean fluorescence intensity (MFI) values confirmed a 1.3-fold increase in fluorescence in the Neu5Ac-supplemented hypersialylated RPE1 group compared to the control, while the desialylated group showed a 0.8-fold decrease in fluorescence (**Figure.2a and b**).

Since sialylation affects cell surface charge, we measured the zeta potential of cells after trypsinization, since measuring the cell surface directly is challenging. Zeta potential is the difference in electric potential at the interface between the mobile aqueous layer and the stationary fluid layer adjacent to the cell surface, called the slipping surface [12], [13]. Post-trypsinization, the zeta potential of the three cell groups was measured in suspension (0.5 × 10^6^ cells/ml) in 1X phosphate-buffered saline (PBS) using a Zetasizer Nano ZS analyzer at 25°C. Control RPE1 cells displayed a zeta potential of -5.98±0.25 mV; hypersialylated cells were significantly more negative at -10.6±2.7 mV, while the desialylated group exhibited a less negative potential of -1.95±0.58 mV (**Figure 2c**). This confirms a change in surface charge across the three treated groups, corresponding to the sialylation levels quantified by WGA staining. The results of WGA staining, along with the zeta potential and viability assay, validate the development of three groups of RPE1 cells with different sialylation states: an untreated control group, a hypersialylated group that mimics the overexpression of sialic acids in cancer cells, and a desialylated group to confirm the contribution of membrane sialylation in TDN uptake.

**Figure 2:**
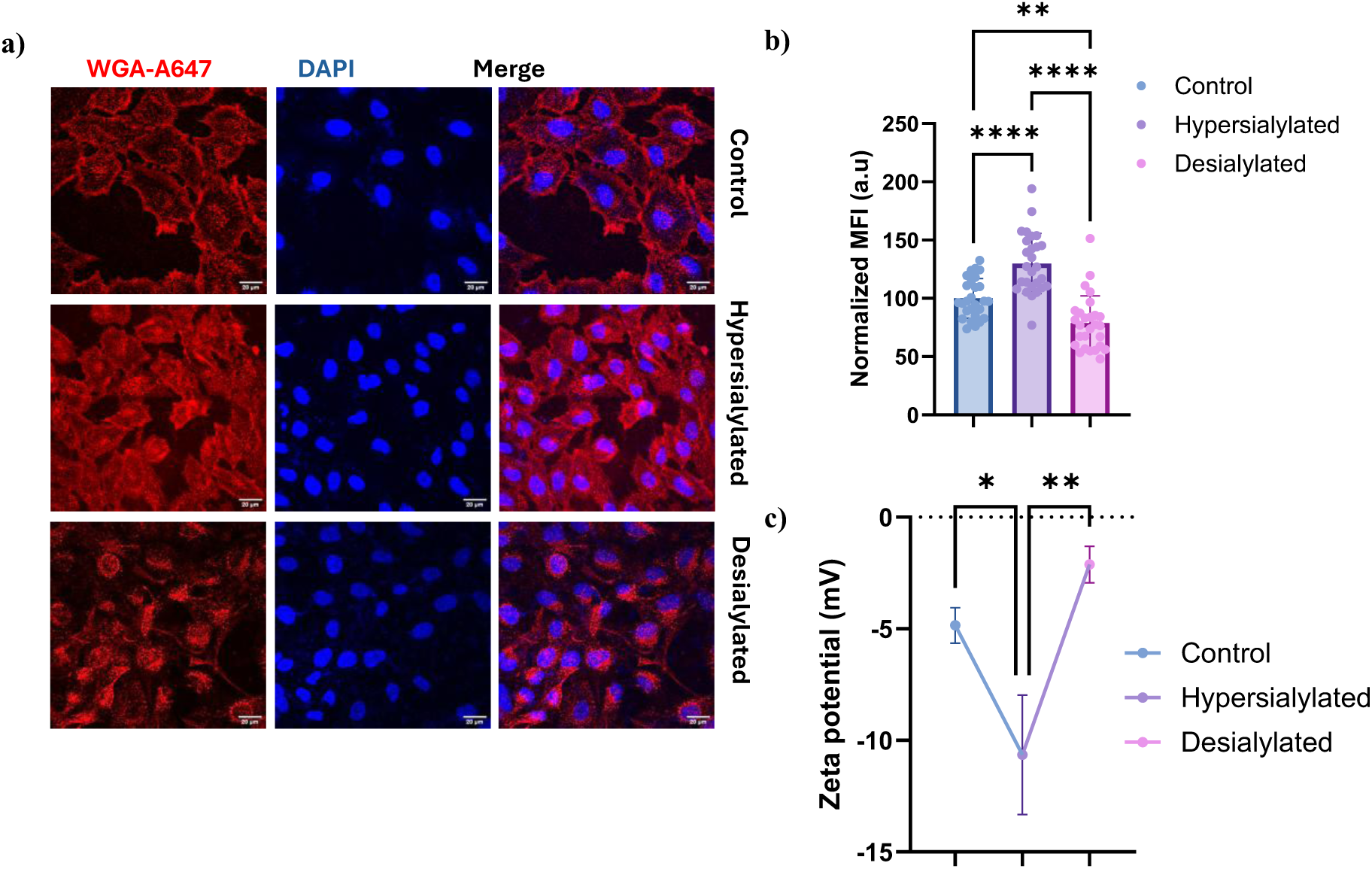
Chemical modulation of sialylation on RPE1 cells. a) Confocal images of WGA staining on the RPE1 models selected for the study. Hypersialylated group was grown on DMEM media supplemented with 2mM Neu5Ac, desialylated group was grown on 2µM 3Fax-Peracetyl Neu5Ac supplemented media, followed by treatment with 0.1U/ml α-2-3,6,8-neuraminidase for 30 minutes, and control cells were grown on normal DMEM growth media. Scale bar is 20µm. b) Quantification of WGA fluorescence showed significant increases and decreases in WGA binding in the hypersialylated and desialylated groups, respectively, compared to the control (Ordinary ANOVA, p-value<0.0001****). Mean± standard deviation is plotted for n=30 cells, N=3. c) Corresponding zeta potential measurement revealed changes in surface charge with altered levels of sialic acids (Ordinary ANOVA, p-value<0.0022** and N=3). Scale bar- 20µm.

### Cell surface sialylation tunes DNA TDN internalization

Using validated RPE1 cell models at different sialylation states, we performed a DNA TDN uptake assay with Cy3-labeled TDN. Cells were incubated with 200nM of Cy3-TDN in serum-free DMEM media for 20 minutes at 37°C, after which the surface-bound TDNs were removed via acid wash (pH 2.5, twice) followed by three washes with 1X PBS [1]. The total intracellular fluorescence was quantified using confocal laser scanning microscopy and flow cytometry. The uptake data revealed a 1.2-fold increase in the hypersialylated group compared to the control, while desialylated cells showed a 2.1-fold increase (**Figure 3.a**). The elevated TDN uptake in desialylated cells is consistent with the expectation that removing anionic sialic acids from the cell surface would reduce electrostatic repulsion of anionic TDN and enhance its interaction with the cell membrane, thereby increasing uptake. However, the excess abundance of anionic sialic acids on hypersialylated cells still led to higher TDN uptake than control cells. This trend contradicts the simple electrostatic model and suggests the involvement of membrane-level mechanisms that override electrostatic repulsion and enhance TDN uptake.

**Figure 3:**
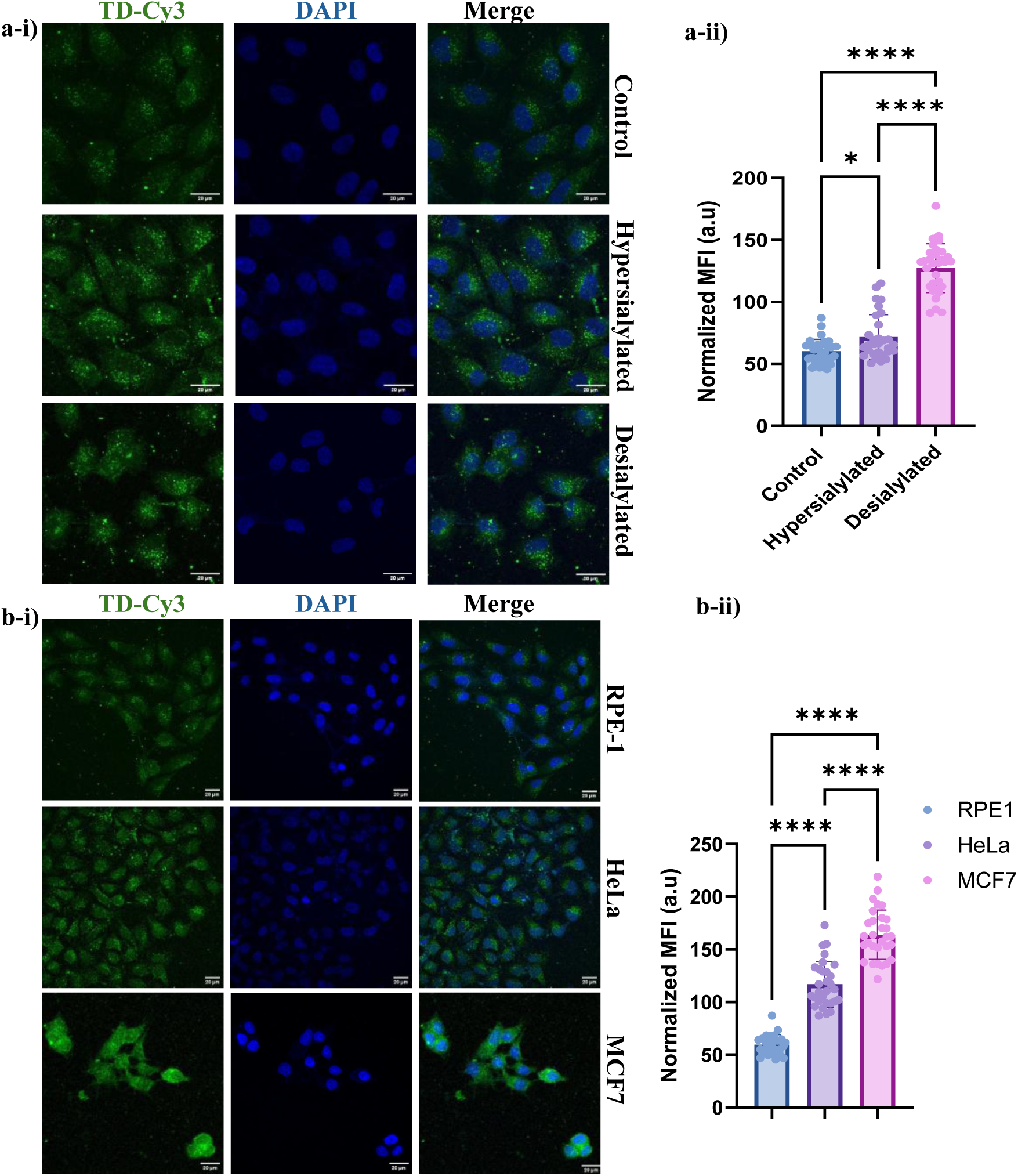
Cellular uptake of TDN under different sialylation states. a-i) Confocal images of cellular uptake of Cy3-TDN (green) in sialylation-modulated RPE1 cells. a-ii) Quantification of fluorescent intensity revealed an increase in TDN uptake upon hyper-sialylation, but a higher increase upon desialylation. b-i) Confocal images of Cy3-TDN uptake in untreated RPE1, HeLa, and MCF7 cells. b-ii) Quantification of TDN fluorescence revealed higher TDN internalization in naturally hypersialylated cancer cells, such as HeLa and MCF7. Data plotted are mean±s.d of n=30 cells. Ordinary one-way ANOVA, p-value> 0.0001**** and N=3. Scale bar is 20µm.

To determine whether the sialylation-driven uptake trend is conserved across naturally hypersialylated human cancer cells (**Figure S5**), we measured TDN uptake in HeLa (cervical carcinoma) and MCF7 (breast adenocarcinoma) cells. Compared with RPE1 cells (control), HeLa showed a 2-fold increase in uptake, while MCF7 showed a 2.8-fold increase (**Figure 3.b**). Although differences in cell lineages and metabolism may contribute to differential TDN uptake between cancerous and non-cancerous cells, the pronounced enhancement in cancer cells, similar to that observed in hypersialylated RPE1 cells mimicking the glycocalyx of cancer cells, provides evidence for the involvement of cell-surface sialic acids in TDN internalization.

### Cell surface sialylation reorganizes lipid rafts to drive DNA TDN uptake

The non-linear pattern of TDN uptake, in which hypersialylated cells still caused higher uptake of TDN despite the higher electrostatic repulsion on the cell surface, requires a mechanistic dissection rather than a purely electrostatic understanding. As studied earlier by Ding et al., negatively charged TDNs overcome repulsion from the negatively charged cell membrane by landing in caveolae containing lipid raft microdomains on the cell membrane, which provide a short-range attractive force that engages the TDN on the cell surface for a longer time, facilitating efficient uptake [14], [15]. This finding led to the understanding that caveolae raft domains at the cell surface are also a determinant of TDN engagement and uptake. Therefore, cells with a greater density of caveolin-recruiting raft domains have more landing sites for TDN and thus greater internalization. This guided us to question whether sialylation levels control the abundance of caveolae rafts on cell membranes. Since GM1 gangliosides, a constituent of the caveolar raft microdomains, are themselves sialylated glycosphingolipids, chemical supplementation of Neu5Ac would increase the biosynthesis of GM1, which subsequently gets inserted into the outer leaflet of the plasma membrane [16]–[19]. This increase in the GM1 pool would expand the GM1-enriched caveolae rafts on the cell membrane, thereby increasing TDN uptake[20], [21].

To answer this question, we employed a staining protocol using the B subunit of Cholera toxin (CTxB) as a specific marker of GM1 microdomains [18], [22], [23]. Quantification of the fluorescent signal from CTxB will therefore report changes in GM1 microdomains and, consequently, changes in caveolae upon increasing membrane sialylation. RPE1 cells were supplemented with increasing concentrations of Neu5Ac (1 mM to 4 mM) in DMEM media for 96 hours, followed by staining using fluorescently labeled CTxB (CTxB-A488). Quantification of fluorescence post confocal microscopy revealed an increase in CTxB fluorescence at 1mM and 2mM Neu5Ac supplementation (**Figure 4.a**), at which WGA staining confirmed a significant increase in membrane sialylation (**Figure S2)**. This finding, therefore, reveals a relation between membrane sialylation and the abundance of GM1-enriched raft microdomains on the cell membrane. To confirm that the increase in CTxB fluorescence upon sialylation truly reflects the abundance of GM1-enriched caveolae recruiting lipid raft microdomains, rather than a simple increase in total surface GM1 density, we employed an inhibition study utilizing Methyl-β-cyclodextrin (MβCD), a member of the cyclodextrin family with high cholesterol-depleting potential. Depletion of cholesterol from the plasma membrane disperses raft-resident caveolin proteins, eliminating caveolae-dependent endocytosis[24], [25]. RPE1 cells grown in Neu5Ac-supplemented media (1mM to 4mM) for 96 hours were subjected to MβCD (10mM) treatment for 15 minutes at 37°C, followed by CTxB staining while maintaining MβCD in media. Quantification of CTxB fluorescence showed a substantial reduction upon MβCD treatment compared with the untreated groups (**Figure S6**). This finding demonstrates that the sialylation-dependent increase in CTxB binding is due to the abundance of cholesterol-dependent, caveolin-containing GM1 raft enrichment at the cell membrane.

**Figure 4:**
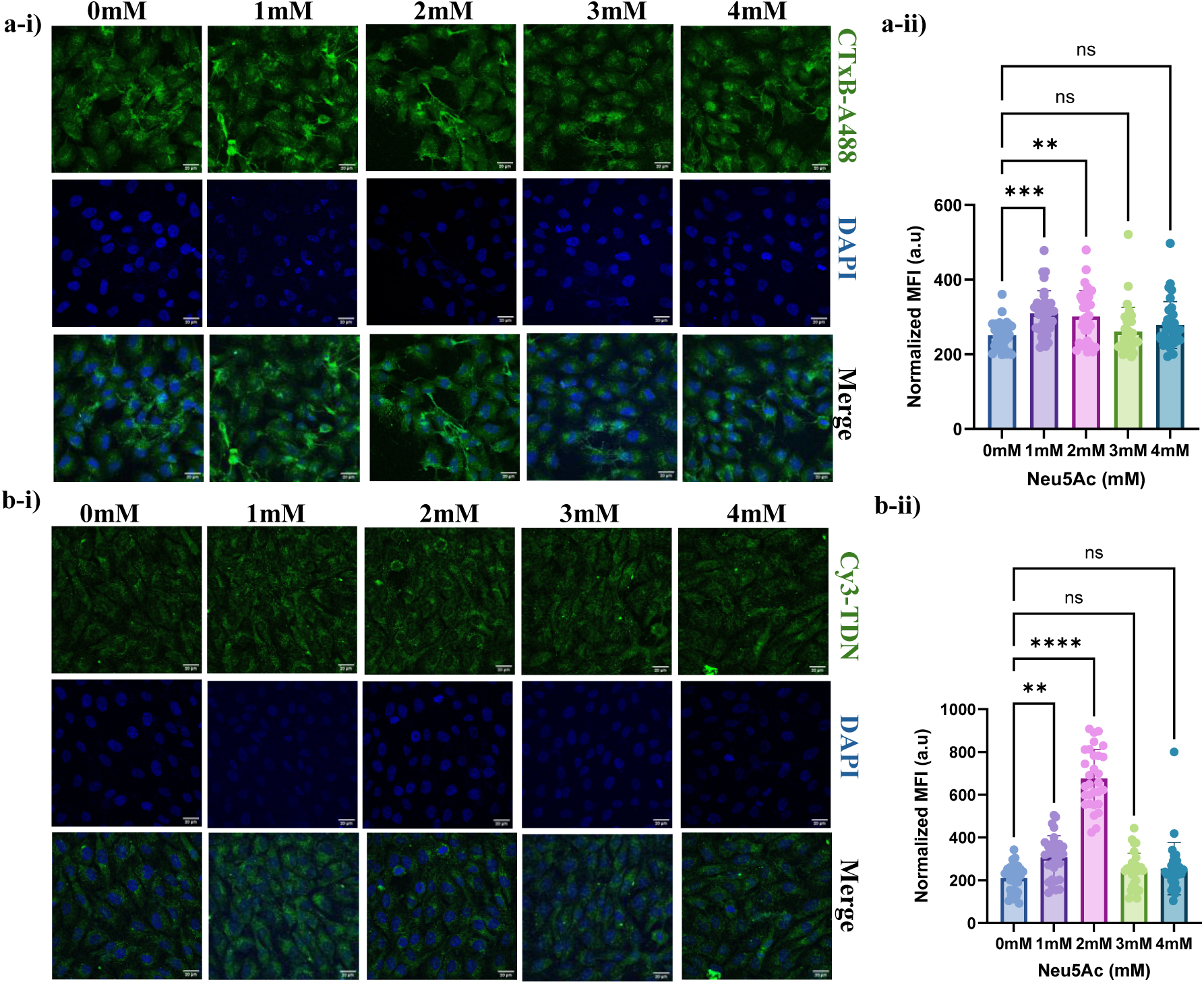
Hyper-sialylation drives alterations in GM1-lipid raft expression on RPE1 cell membrane and enhances TDN uptake. a-i) Confocal images of CTxB staining in RPE1 cells grown in increasing concentrations of Neu5Ac (0mM-4mM). a-ii) Quantification of fluorescence intensity shows an increase in CTxB binding at 1 mM and 2 mM Neu5Ac treatment, indicating an abundance of the GM1 microdomain. Data plotted are mean±s.d, n=30 cells, N=3, Ordinary one-way ANOVA, and p-value < 0.0002***). b-i) Confocal images of Cy3-TDN uptake in RPE1 cells grown at similar concentrations of Neu5Ac. b-ii) Quantification of CY3-TDN fluorescence revealed a significant increase at the same Neu5Ac concentrations (1 mM and 2 mM). Data plotted are mean±s.d, n=30 cells, N=3, one-way ANOVA, and p-value < 0.0001****). Scale bar- 20µm.

With the understanding that hyper-sialylation quantitatively increases the number of caveolae-containing raft microdomains at the cell surface, we directly quantified TDN uptake to test whether the sialylation-driven enhancement of caveolae-containing rafts increases TDN uptake. If so, MβCD-driven disruption of caveolae at the cell membrane should reduce TDN uptake in Neu5AC-treated cells compared to control groups not supplemented with Neu5Ac, where raft-dependent uptake is expected to be comparatively lower. Cy3-labeled TDN were incubated in the presence and absence of MβCD with RPE1 cells grown for 96 hours in Neu5Ac-supplemented (1mM to 5mM) media. Quantification of TDN fluorescence revealed a significant increase up to 1mM and 2 mM Neu5Ac, in the absence of MβCD, similar to the CTxB profile observed earlier (**Figure 4.b**). However, upon MβCD treatment, TDN uptake was significantly reduced at the Neu5Ac concentrations (**Figure S7**). These findings demonstrate that the increase in TDN uptake upon hyper-sialylation is driven by the abundance of caveolae containing lipid-raft microdomains on the cell membrane.

### Alterations in cell membrane sialylation switch the endocytic pathway of TDN uptake

Having understood the abundance of caveolae during hyper-sialylation and the subsequent increase in TDN uptake, we sought to dissect the endocytic pathway operating under each condition. We employed a set of chemical inhibitors specific to three principal pathways of endocytosis: Clathrin-mediated endocytosis (CME), Caveolae-mediated endocytosis (Cav), and Clathrin-independent endocytosis (CIE), and quantified the TDN uptake in each condition. Table 1.

**Table 1:**
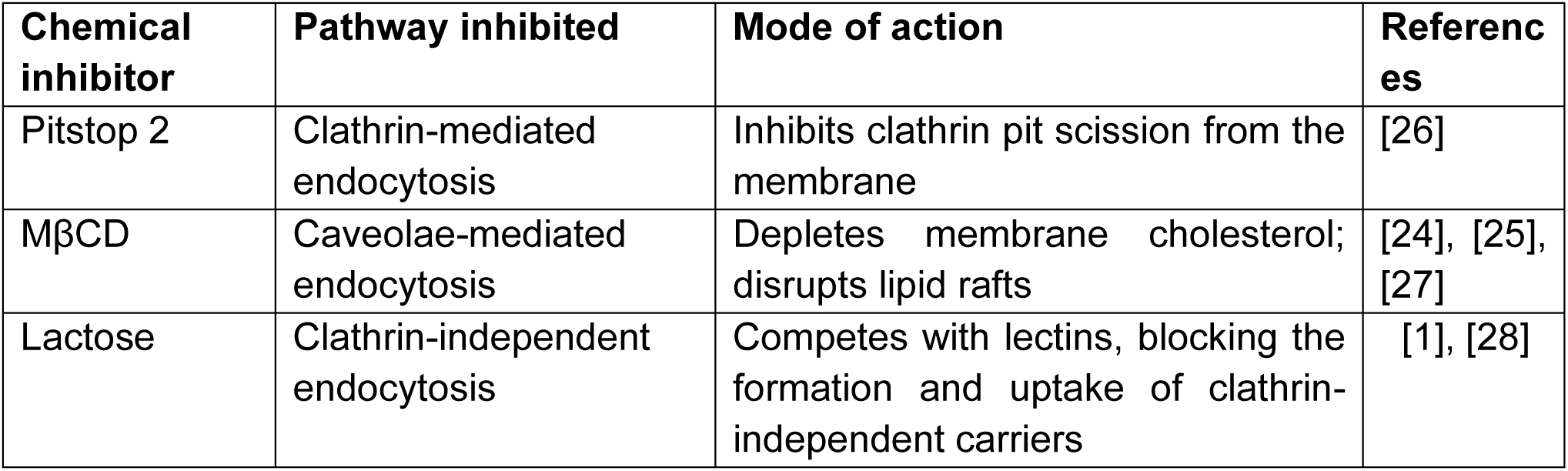
Chemical inhibitors used to dissect endocytic pathways mediating TDN uptake in sialylation-altered RPE1 cells.

| Chemical inhibitor | Pathway inhibited | Mode of action | References |
| --- | --- | --- | --- |
| Pitstop 2 | Clathrin-mediated endocytosis | Inhibits clathrin pit scission from the membrane | [26] |
| MβCD | Caveolae-mediated endocytosis | Depletes membrane cholesterol; disrupts lipid rafts | [24], [25], [27] |
| Lactose | Clathrin-independent endocytosis | Competes with lectins, blocking the formation and uptake of clathrin-independent carriers | [1], [28] |

In untreated RPE1 cells expressing baseline sialylation, Pitstop-2 treatment reduced TDN uptake by 60%, while MβCD reduced it by 10%, and lactose had no significant effect, confirming CME as the principal endocytic pathway of TDN uptake under baseline conditions, with minor contributions from Cav, while CIE played no role (**Figure 5.a**). However, in hypersialylated RPE1 cells, TDN uptake showed a greater reduction in the presence of MβCD, up to 40%, while Pitstop-2 reduced uptake by 60%, and CIE remained ineffective. This reduction in TDN uptake in the presence of both Pitstop-2 and MβCD indicates the involvement of both pathways during hyper-sialylation, thereby enhancing TDN uptake compared to baseline conditions (**Figure 5.b**). In desialylated cells, Pitstop-2 reduced uptake by 46%, and no significant reduction in the presence of MβCD or lactose. This indicates a complete reduction in Cav involvement during desialylation (**Figure 5.c**). Taken together, these data reveal that cell surface sialylation is a determinant of the endocytic route for TDN uptake, with hyper-sialylation significantly increasing both CME and Cav involvement. Importantly, this shift towards the caveolar pathway for TDN internalization under hyper-sialylation provides a mechanistic basis for understanding why caveolin-mediated endocytosis becomes a major pathway of TDN uptake in intrinsically hypersialylated cancer cells like HeLa [1], [15], [29], [30] .

**Figure 5:**
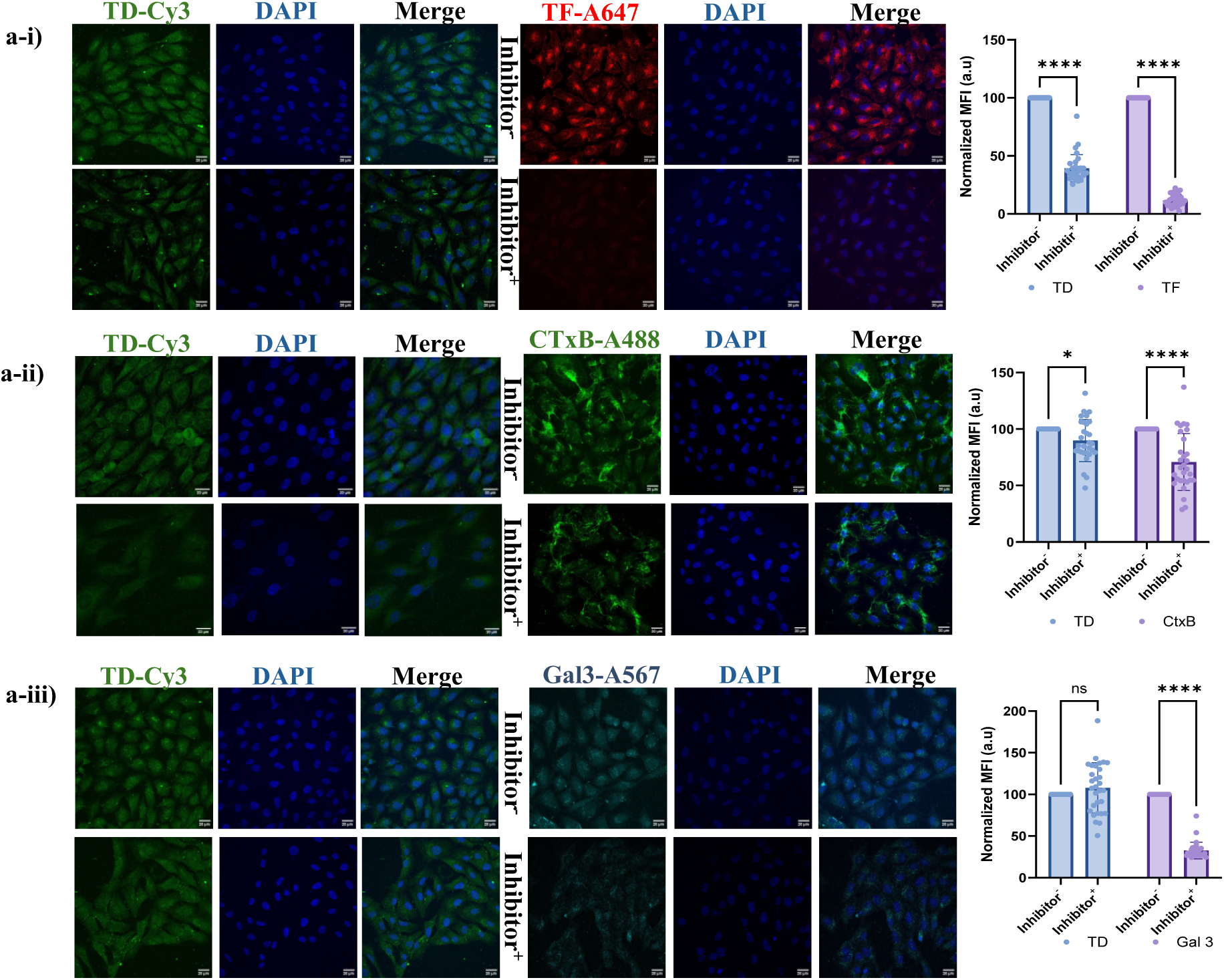
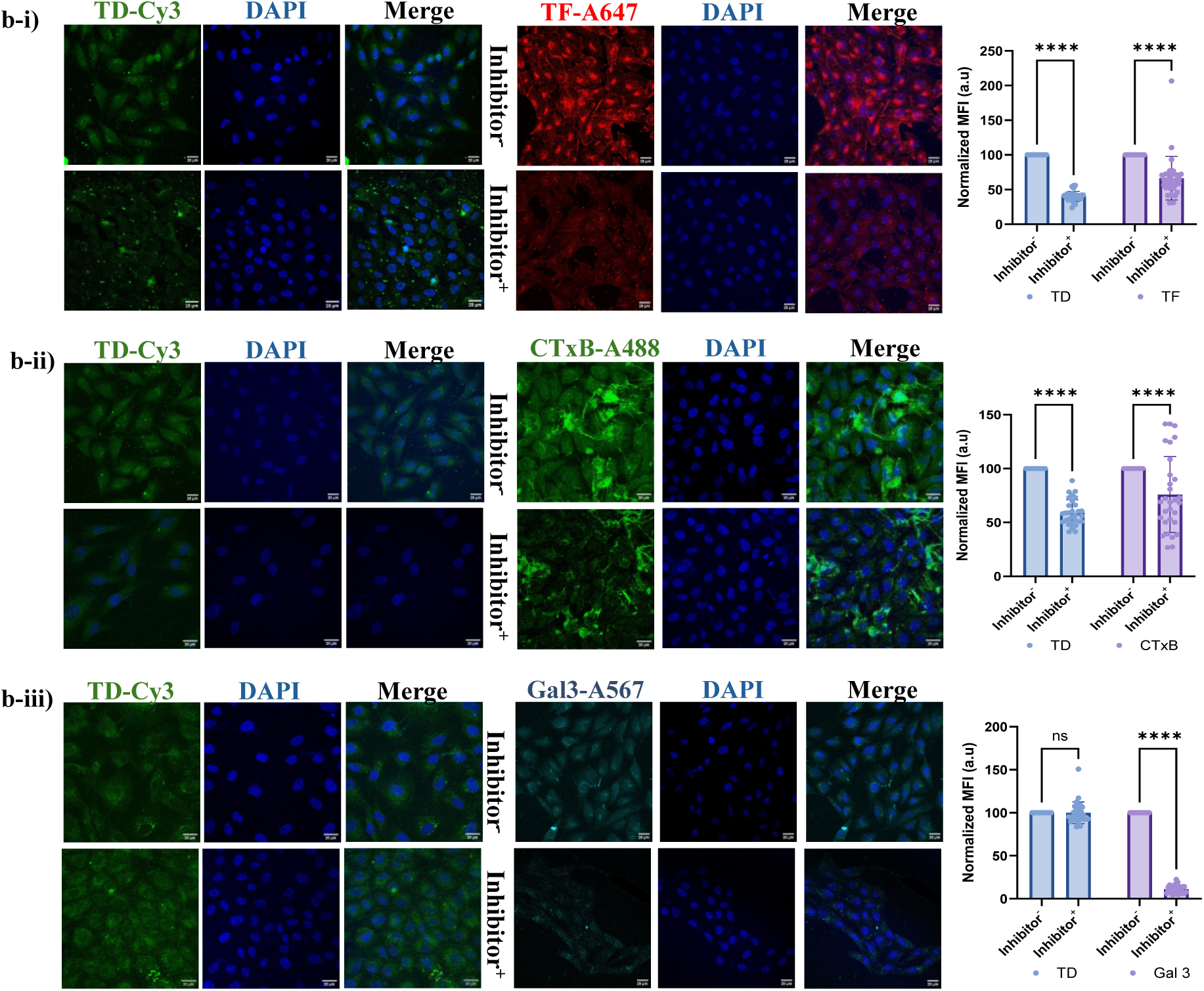
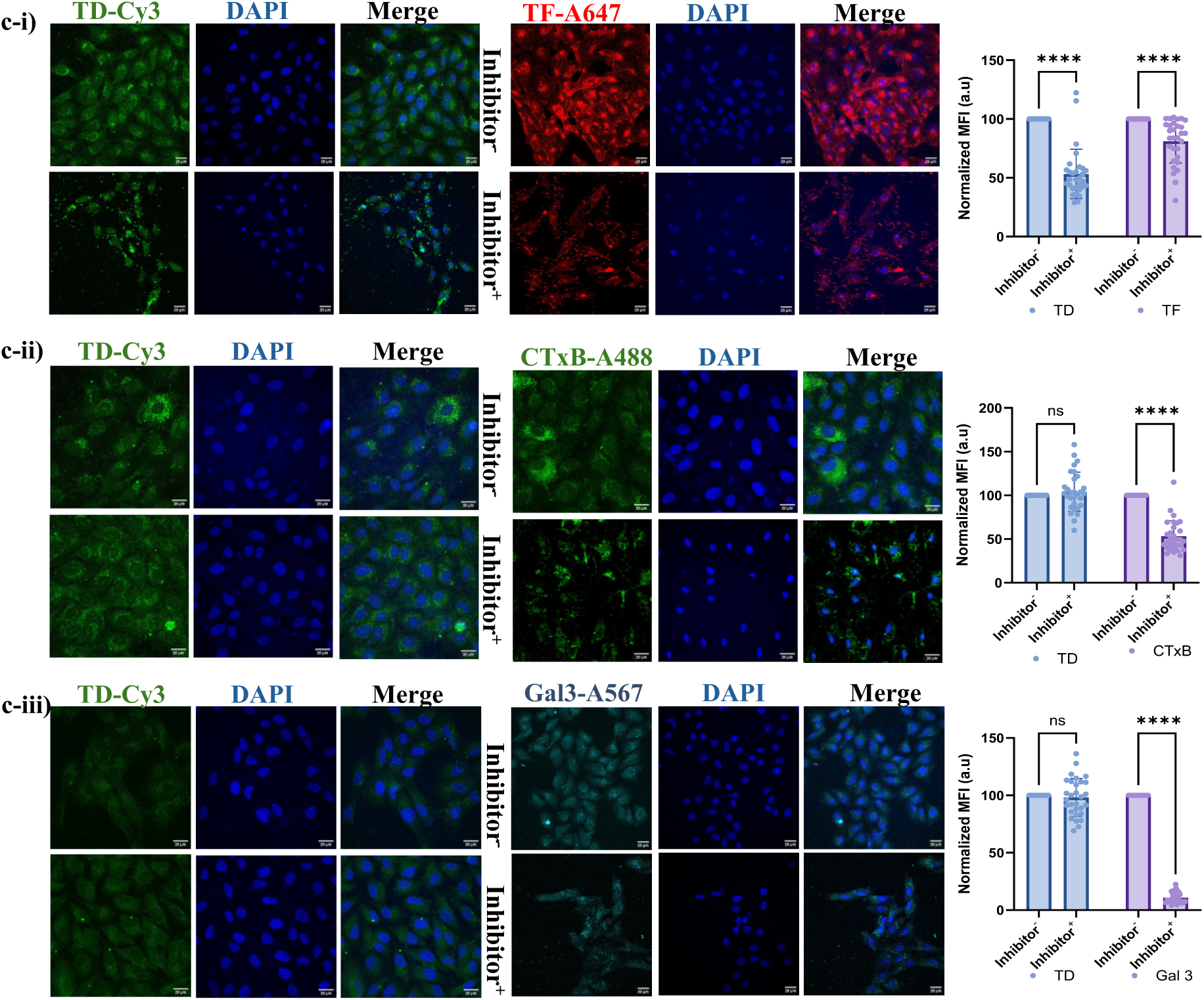
Membrane sialylation modulates the TDN uptake pathway. Confocal images dissecting DNA TDN (green) uptake via chemical inhibitor-based inhibition of endocytic pathways. a) RPE1 cells expressing baseline sialylation (control) showed a significant reduction in TDN uptake in the presence of CME inhibitor, Pitstop-2 (a-i), while in the presence of Cav inhibitor (MβCD), the uptake was mildly reduced (a-ii); however, no reduction was observed in the presence of CIE inhibitor (lactose, a-iii). b) Hypersialylated RPE1 cells showed a significant reduction in TDN uptake in the presence of both Pitstop-2 (b-i) and MβCD (b-ii), with no reduction in the presence of lactose (b-iii). This indicates equal involvement of both CME and Cav in TDN uptake under hyper-sialylation. c) Desialylated RPE1 cells showed a significant reduction in uptake in the presence of Pitstop-2 (c-i), while no reduction was observed in the presence of MβCD (c-ii) or lactose (c-iii), indicating the involvement of CME only. Inhibition was confirmed in each case using validated cargoes for the respective pathways: transferrin (TF, red) for CME, CTxB (green) for Cav, and galectin-3 (Gal-3, Cyan) for CIE. Data plotted are mean±s.d, n=30 cells, N=4, Two-way ANOVA, and p-value < 0.0001****). Scale bar- 20µm.

### Sialylation-dependent uptake pattern extends to 3D spheroid models

To determine whether the sialylation-dependent uptake of TDN pattern observed in 2-dimensional cell monolayers extends to physiologically complex tumor models, we performed a TDN uptake assay in 3D spheroid models generated from each of the glycoengineered RPE1 cells using the hanging drop method [31]. Suspensions of RPE1 cells grown under the above-discussed conditions for 96 hours were placed as 20 µL drops on the lid of a petri dish containing 1x PBS. The cells aggregated under gravity and formed compact spheroids within 36 hours, which were later transferred to a collagen matrix on coverslips. The spheroids were incubated with 200nM of Cy5-labeled TDN for 24 hours at 37°C. Confocal microscopy revealed a higher TDN uptake (1.3-fold) in hypersialylated spheroids compared to the control group, while desialylated spheroids showed a 1.7-fold increase (**Figure 6**).

**Figure 6:**
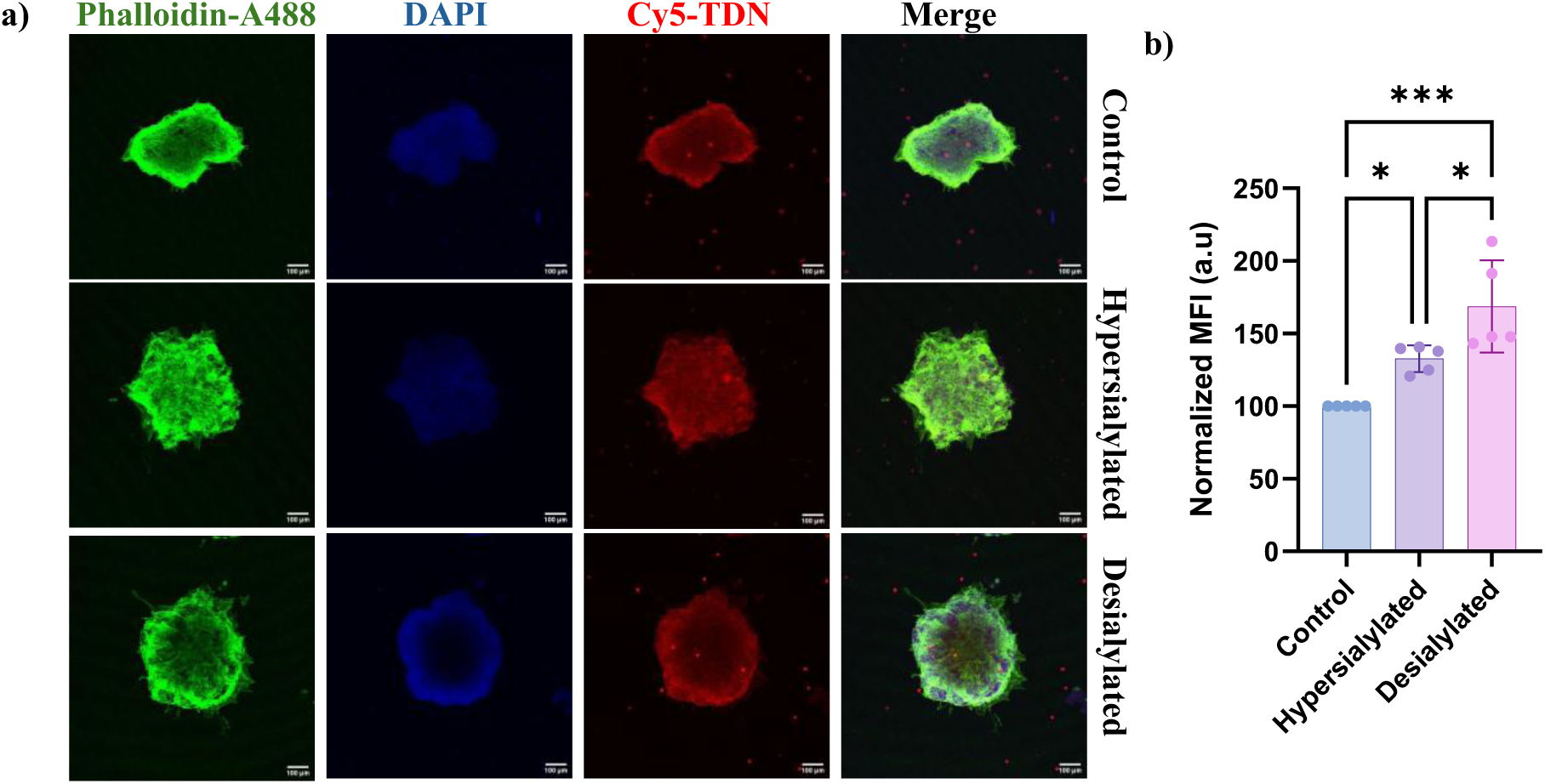
Sialylation-dependent uptake of TDN extends to 3D spheroids. a) Confocal images of Cy5-labeled TDN (red) uptake in sialylation-modulated RPE1 spheroids. Phalloidin staining (green) was performed to visualize the actin filaments, along with DAPI to visualize nuclei (blue). b) Quantification of TDN fluorescent intensity. A similar trend as observed in 2D RPE1 monolayers was observed. Data represent mean±s.d. 5 spheroids were studied per condition. (One-way ANOVA with P<0.0004***) and N=3, scale bar in a is 100μm.

### In vivo validation in zebrafish confirms sialylation-dependent uptake of DNA TDN

To provide in vivo validation of the sialylation-dependent uptake of TDN, we employed zebrafish (*Danio rerio*) embryo models, a promising tool for screening nanoparticle uptake [32]. The sialylation levels of zebrafish embryos were altered to establish a hypersialylated and desialylated model to validate our findings. Hypersialylated models were developed by growing zebrafish embryos in E3 media supplemented with increasing concentrations of Neu5Ac (0.01 mM to 0.05 mM) for 96 hours post-fertilization (hpf), as determined by a lethal concentration 50 (LC50) toxicity assessment. Based on the LC50 values, all working concentrations were set at 1/10 of the LC50 (0.5mM) concentration to ensure complete viability and the absence of developmental toxicity (**Figure S8**). Similarly, desialylated models were developed by growing embryos in 3Fax-Peracetyl Neu5Ac (1 µM to 4 µM) for 96 hpf. Sialylation level was confirmed via WGA staining after 96 hours of treatment, followed by confocal imaging of whole larvae (**Figure S9**) [33]. Quantification of WGA fluorescence indicated an increase in sialylation level at 0.05mM Neu5Ac, while at 1µM supplementation of 3Fax-Peracetyl Neu5Ac, membrane sialylation significantly reduced (**Figure 7a**). To study TDN uptake, Cy5-labeled TDN was used and fed in E3 media at a 200nM concentration for 4 hours [1], [34]. Confocal imaging revealed an increase in TDN fluorescence at 0.05mM Neu5Ac (**Figure 7b**).

**Figure 7:**
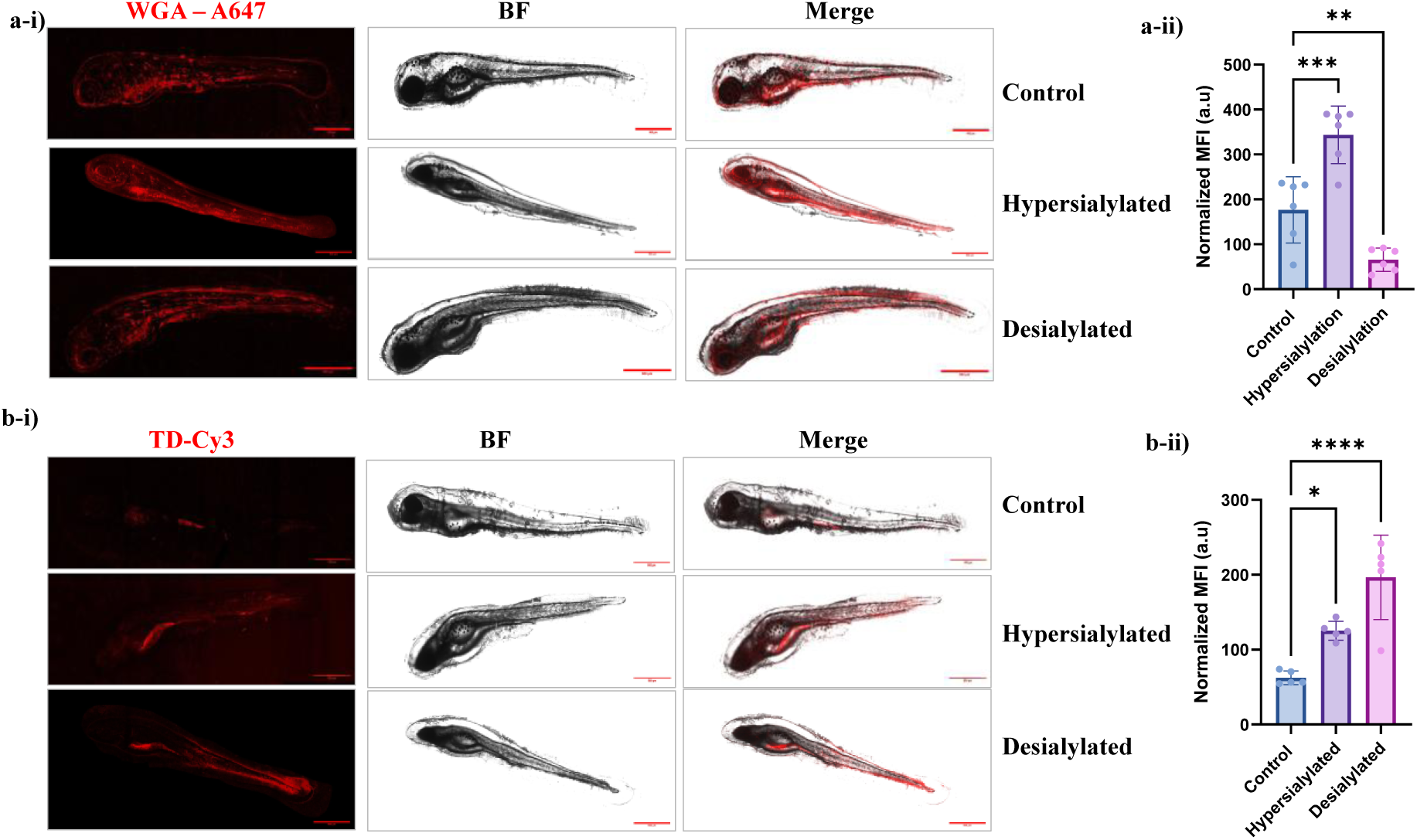
In vivo validation of sialylation-dependent TDN uptake in zebrafish. a-i) Confocal images of WGA staining on zebra fish larvae at 96hpf, grown in E3 media (control), E3 media supplemented with 0.05mM Neu5Ac (hypersialylated), and E3 media supplemented with 1µM 3Fax-Peracetyl Neu5Ac. a-ii) Quantification of WGA staining showed higher WGA binding in hypersialylated larvae. p-value<0.0001****. b-i) Confocal images of Cy5-TDN uptake in zebrafish larvae (96hpf) with varying sialylation. b-ii) Quantification of TDN fluorescence revealed increased TDN uptake in the hypersialylated group compared to the control group, whereas the desialylated group showed the highest significant TDN uptake. p-value<0.0002***. Data presented is mean±s.d. 6 larvae were counted per treatment, N=3. Scale bar is 500µm.

Taken together, our findings provide insight into how anionic DNA nanostructures, such as TDN, interact with the cancer cell surface. Rather than a passive electrostatic barrier, cell-surface sialylation actively modulates the internalization of these nanostructures by expanding the GM1 raft microdomains that contain caveolin. This reveals how the overexpression of sialic acids, a hallmark of cancer cells, enables efficient internalization of DNA nanostructures.

## Conclusion

We have demonstrated that cell-surface sialylation is a quantitative and mechanistic determinant of DNA TDN uptake, operating through the coupled modulation of electrostatic forces and lipid raft architecture at the cell membrane. This coupling produces a sialylation-dependent switch in the endocytic pathway for TDN uptake, with CME as the principal pathway under normal conditions, along with a quantitatively lesser contribution from caveolae. However, upon hyper-sialylation, caveolae also play a prominent role as an additional pathway for TDN uptake, thereby increasing total TDN internalization. In the absence of sialic acids, caveolar involvement was inhibited, leaving CME as the sole mechanism of uptake. These findings establish the membrane glycocalyx as a driver of nanostructure uptake and suggest that the hypersialylated tumor glycocalyx, a molecular signature that cancer cells use for immune evasion, simultaneously expands the caveolae raft architecture, which facilitates the entry of anionic DNA nanostructures. The principle that surface sialylation encodes endocytic pathway identity provides a new conceptual framework for the rational design and delivery optimization of DNA-based nanostructures, with direct implications for cancer targeting.

## Methods

### 1. Materials

The oligonucleotide sequence used for TDN synthesis (**Table S1**) was purchased from Sigma. Nuclease-free water (SRL), magnesium chloride hexahydrate (Merck), 30% acrylamide/bis-acrylamide (29:1) (SRL), 6X gel loading dye (Promega), ethidium bromide (Himedia), High-glucose Dulbecco’s Modified Eagle’s Medium (Gibco), fetal bovine serum (Gibco), 0.25% trypsin-EDTA (Gibco), N-Acetylneuraminic acid (Carbosynth), 3Fax-Peracetyl Neu5Ac (Calbiochem), α-2-3,6,8-neuraminidase (New England Biolabs), Wheat germ agglutinin–Alexa Fluor 647 (Invitrogen), cholera toxin subunit B–Alexa Fluor 488 (Invitrogen), transferrin–Alexa Fluor 647 (Invitrogen), phalloidin–Alexa Fluor 488 (Invitrogen), DAPI (Sigma-Aldrich), Pitstop-2, methyl-β-cyclodextrin (Sigma-Aldrich), lactose (Fisher Scientific), Paraformaldehyde (SRL), Triton X-100 (Himedia), Collagen type I from rat tail (Corning), cell-culture dishes (NEST). Galectin-3 was provided as a gift from the Johannes team at Institut Curie, Paris.

### 2. Synthesis and characterization of DNA TDN

#### 2.1 Synthesis of TDN

To synthesize DNA TDN, we used a one-pot synthesis strategy in which four single-stranded complementary DNA primers were reconstituted in nuclease-free water to a final working concentration of 10 µM each, in equimolar ratio, in the presence of 2 mM MgCl_2_. TDN synthesis was carried out in a T100 PCR Thermal Cycler with an initial denaturation stage at 95°C for 30 minutes, followed by a 5 °C reduction with a 15min incubation at each step, until 4 °C. The final concentration of TDN was 2.5µM. For fluorescence-based uptake experiments, one primer was used with 5’-Cy3 or 5’-Cy5.

#### 2.2 Characterization of TDN

##### a) Electrophoretic mobility shift assay (EMSA)

The formation of tetrahedral structure was primarily confirmed by Electrophoretic mobility shift assay (EMSA) using 10% native polyacrylamide gel (PAGE). 5µl of TDN with 3µl of 1X TAE buffer and 1.5 µl of 6X loading dye was used. Electrophoresis was carried out at 80V for 90 minutes, followed by staining of the gel with 0.32 µg/ml ethidium bromide prepared in 1X TAE buffer for 10 minutes, and visualized using the Bio-Rad ChemiDoc MP gel documentation system.

##### b) Dynamic light scattering (DLS)

Hydrodynamic diameter and zeta potential of TDN were measured using Zetasizer Nano ZS (Malvern Panalytical) at 25°C in nuclease-free water. For each measurement, three independent runs were performed.

##### c) Atomic force microscopy (AFM)

The morphology of the synthesized TDN was characterized using AFM. The TDN sample was diluted 10X in nuclease-free water, drop-casted onto a fresh mica sheet, and vacuum-dried. Once dried, TDN was analyzed using an Atomic Force Microscope (Bruker; Nano wizard) in air tapping mode (AFM probe: Peakforce HIRS-F-A, tip radius: 1nm, frequency: 165 kHz, line rate: 0.4 lines/sec). The images were analyzed using JPK data processing software.

### 3. Cell Culture

Cells utilized in the study were grown under standard incubation conditions of 37°C and 5% CO_2_ in a humidified chamber. RPE1, HeLa, and MCF7 were grown in DMEM high-glucose (4.5 g/L) media supplemented with 10% FBS and 1% penicillin-streptomycin (complete media).

#### 3.1 Chemical glycoengineering of RPE1 cells

##### a) Hyper sialylation

RPE1 cells grown in DMEM complete media were passaged at 80% confluency using trypsin-EDTA (0.25%) and seeded at a 10,000 cells/well density on coverslips placed on 24- well cell culture plates. Neu5Ac stock solution was prepared at a 100 mM concentration in 1X PBS. Final Neu5Ac concentrations ranging from 1 mM to 4 mM were prepared in DMEM complete media and used during seeding[35], [36]. Cells were grown in Neu5Ac-supplemented media for 96 hours, with media replacement every 48 hours.

##### b) Desialylation

To achieve near-complete removal of sialic acids from RPE1 cells, the DMEM complete media was supplemented with 3Fax-Peracetyl Neu5Ac (1 µM to 3 µM) for 96 hours on cells seeded as mentioned, and the media was replaced every 48 hours [8]. This was followed by a 30-minute incubation with 0.1U/ml of α-2-3,6,8-neuraminidase at 37°C.

##### c) Control

Untreated RPE1 cells grown in normal DMEM complete media served as the control with baseline sialylation.

###### i) Wheat germ agglutinin (WGA) staining

Cell surface sialylation was quantified using WGA-Alexa 647 at a working concentration of 5µg/ml in DMEM serum-free media. Cells grown in the presence of treatments to alter sialylation were subjected to WGA staining at 96 hours. Cells were washed twice with 1X PBS to remove any debris, then incubated with WGA-A647 prepared in serum-free media for 10 minutes at 37°C and 5% CO_2_. Post-incubation, the cells were washed 3 times with 1X PBS to remove unbound lectins, followed by fixation with 4% paraformaldehyde (PFA) prepared in 1X PBS for 15 minutes at 37°C and 5% CO_2._ The cells were later counterstained with DAPI for nuclear visualization and imaged by confocal microscopy.

###### ii) Cell-surface zeta potential measurement

RPE1 cells grown on different treatment conditions were subjected to trypsinization (0.25% trypsin-EDTA, for 5minutes at 37°C and 5% CO_2_) to create a cell suspension. The cells were pelleted and washed with 1X PBS. Cell counting was performed using a hemocytometer, and equal concentrations of 0.5 × 10^6^ cells/ml were maintained in each condition. Zeta potential was measured at 25°C in U-shaped cuvettes containing gold-plated electrodes using Zetasizer Nano ZS [13], [37], [38].

#### 3.2 Cell uptake assay

To study the cellular uptake of DNA TDN in sialylation-modulated RPE1 cells, Cy3-labeled TDN was used. Cells were grown on coverslips in a 24-well plate at a density of 10,000 cells/well under the specified treatment conditions for sialylation modulation, as described above for 96 hours. Cells were observed daily under a microscope for attachment and morphology. After 96 hours of incubation, the cells were washed twice with 1X PBS, then incubated with 200nM Cy3-TDN prepared in DMEM serum-free media for 20 minutes at 37°C and 5% CO_2._ The cells were washed twice with acid buffer (0.2 M acetic acid + 0.5 M NaCl at pH 2.5) to remove surface-bound TDN, followed by three washes with 1X PBS. The cells were then fixed using 4% paraformaldehyde for 15 minutes at 37°C. After two washes with 1X PBS, the coverslips containing cells were mounted on glass slides using Mowiol and DAPI as a counterstain to stain the nucleus. The fluorescence of TDN was measured using a confocal laser-scanning microscope with a 63X oil-immersion objective.

The same protocol was used for the uptake assay performed on HeLa and MCF7 cells, except for the treatment conditions used to modulate sialylation. HeLa and MCF7 cells were seeded and maintained in DMEM complete media for the experiment.

After confocal imaging, the fold increase in uptake was calculated as:

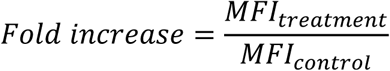

#### 3.3 Lipid raft quantification

##### 3.3.1 CTxB staining to quantify lipid raft abundance

To quantify the changes in the raft microdomain, CTxB staining was used. RPE1 cells were seeded on coverslips of a 24-well plate at 10,000 cells/well density. The cells were grown in DMEM complete media supplemented with increasing concentrations of Neu5AC (1 mM to 5 mM) for 96 hours. After 96 hours, CTxB staining was performed in DMEM serum-free media at a final working concentration of 0.5 µg/ml for 30 minutes at 37°C and 5% CO2, following two 1X PBS washes. Cells were washed twice after CTxB staining, then fixed with 4% paraformaldehyde. Coverslips containing cells were mounted on glass slides using Mowiol, and nuclei were stained using DAPI.

##### 3.3.2 MβCD-based raft disruption study

To confirm that the elevated signal of CTxB is due to raft abundance, we used MβCD, a cholesterol-depleting agent that, in turn, disrupts the caveolae-containing membrane raft microdomain. RPE1 cells seeded on coverslips and grown on Neu5Ac supplementation for 96 hours, as mentioned above, were subjected to incubation with MβCD at 10mM prepared in serum-free media for 15 minutes at 37°C. This was followed by incubation with CTxB as described above in DMEM serum-free media containing 10mM MβCD. After incubation, the cells were washed with 1X PBS and fixed with 4% paraformaldehyde.

Cell uptake assay of Cy3-labeled TDN was performed as discussed above at similar Neu5Ac concentrations (1 mM to 5 mM), along with MβCD-based inhibition. The TDN concentration (200nM) and incubation conditions were kept constant in both cases, as mentioned previously in (**3.2 Cell uptake assay**).

#### 3.4 Inhibitor-based study of endocytic pathways

Chemical inhibitors were used to study the endocytic pathway in RPE1 cells during altered membrane sialylation. RPE1 cells seeded (10,000 cells/well) on coverslips of 24-well plates were grown under different treatments (2mM Neu5Ac for hyper sialylation, 2µM 3Fax-Peracetyl Neu5Ac with 1U/ml of neuraminidase treatment for desialylation, and untreated control). After 96 hours of incubation, the cells were washed twice with 1X PBS and then preincubated with respective inhibitors. Pitstop-2 (20 µM), MβCD (10mM), and lactose (100 mM) in DMEM serum-free media were utilized to inhibit CME, Cav, and CIE, respectively. The cells were exposed to the inhibitors for 15 minutes at 37°C. Inhibition was confirmed using fluorescently labeled, validated cargo for each pathway: TF for CME, CTxB for Cav, and Gal-3 for CIE. Post-inhibitor exposure, the cells were treated with DMEM serum-free media containing the same concentrations of inhibitors, along with TF-A647 (5 µg/ml), CTxB-A488 (0.5 µg/ml), Galectin-3-A567 (5 µg/ml), and Cy3-TDN (200nM) for 20 minutes at 37°C. The cells were then washed with 1XPBS, fixed with 4% paraformaldehyde, and mounted on coverslips using Mowiol, with DAPI to stain nuclei. Confocal imaging with a 63X oil-immersion objective was used to quantify uptake.

Percentage inhibition was calculated as:

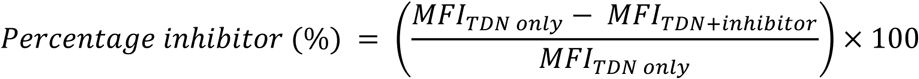

#### 3.5 Uptake assay in 3D spheroid model

RPE1 cells were grown under three conditions in a 6-well plate: control cells grown in DMEM complete media, hypersialylated cells grown in 2mM Neu5Ac-supplemented DMEM complete media, and desialylated cells grown in the presence of 2µM 3Fax-Peracetyl Neu5Ac, to alter membrane sialylation for 96 hours. After 96 hours, the cells were trypsinized and pelleted. 0.5×10^6^ cells were suspended in 1ml of fresh DMEM complete media along with the respective treatments. 20 µL of cell suspension was placed as droplets on the lid of a Petri plate, and the base of the Petri plate was filled with 15 ml of 1X PBS to create a humidified environment for spheroid development via the hanging drop method, in which spheroids form in the hanging drop due to gravity at 37°C and 5% CO_2_. After 36 hours, the spheroids were transferred to coverslips coated with 50 µL of collagen-DMEM complete media (3:1 vol/vol) and incubated for 1 hour at 37°C. The spheroids were then incubated with Cy5-labeled TDN in DMEM complete media for 24 hours at 37°C. Spheroids were then fixed with 4% paraformaldehyde for 15 minutes at 37°C, followed by permeabilization with 0.1% Triton-X for 10 minutes. Post-permeabilization, spheroids were incubated in a solution of 0.1% Triton-X and Phalloidin-A488 (1:1000 dilution) for 30 minutes at 37°C to stain actin filaments. After staining, the spheroids were gently washed with 1X PBS, then mounted with Mowiol and DAPI to stain nuclei. The spheroids were visualized using confocal microscopy with a 10X objective.

### 4 In-Vivo studies

#### 4.1 *Danio rerio* (Zebrafish) husbandry and maintenance

Adult wild-type zebrafish (*Danio rerio*) were maintained in a recirculating zebrafish housing system (GENDANIO BIOTECH INC, Taiwan). Fish were housed under standard laboratory conditions described in ZFIN (Zebrafish Information Network), at 28 ± 0.5°C with a 14:10 h light: dark cycle. The system water was maintained at pH 7.0–7.5, conductivity of ∼300-350 µS/cm, and dissolved oxygen levels above 6 mg/L. The fish were fed twice daily with a combination of commercially available dry feed and live food (e.g., *Artemia* nauplii). Embryos were obtained through natural pairwise mating (3 females: 2 males) and collected within 1-hour post-fertilization (hpf). Embryos were raised in E3 embryo medium (5 mM NaCl, 0.17 mM KCl, 0.33 mM CaCl2, and 0.33 mM MgSO_4_) at 28.5°C and staged according to standard developmental criteria.

##### 4.1.1 Toxicity assessment

Embryos were monitored daily for survival and for gross morphological abnormalities, including body curvature, cardiac edema, and bladder inflation, using a stereomicroscope.

#### 4.2 Chemical glycoengineering in zebrafish

Chemical glycoengineering of zebrafish was performed by growing embryos from 1hpf to 96hpf at 28°C in E3 media supplemented with the treatments listed below. To induce hyper-sialylation, the embryos were grown in E3 media supplemented with 0.05 mM Neu5Ac, and for desialylation, we used 1 µM of 3Fax-Peracetyl Neu5Ac, while embryos grown in E3 media served as the baseline control. The concentrations were optimized following the LC50 toxicity assessment in embryos. The embryos were observed every 24 hours under an upright light microscope for morphology. Changes in sialylation levels in treated groups were validated using WGA staining.

##### 4.2.1 WGA staining

Embryos grown for 96hpf under treatment conditions were subjected to WGA staining by incubating the larvae at 96hpf in E3 media containing 1µg/ml WGA-Alexa 647 for 10 minutes at 28°C. After staining, the larvae were washed 3 times in E3 media, fixed with 4% paraformaldehyde for 2-3 minutes, washed with E3 media, and mounted on a glass slide using a viscosity-enhanced mounting medium. Quantification of WGA fluorescence was performed using confocal microscopy with a 10X objective.

#### 4.3 TDN uptake assay

Zebrafish larvae at 96hpf grown on respective treatments were incubated with Cy5-labeled TDN prepared in E3 media at 200nM concentration for 4 hours at 28°C. The treatment was carried out on 6-well plates, with 10 larvae per condition. Post-incubation, the larvae were washed twice with E3 media, then fixed with 4% paraformaldehyde for 2-3 minutes, washed 3 times with E3 media, and mounted on a glass slide using a viscosity enhanced-mounting medium. Three independent experiments were performed with 6 larvae per treatment.

### 5 Statistical analysis

All data are represented as mean± standard deviation (S.D) from a minimum of three independent biological replicates. Statistical comparisons were performed using one-way ANOVA for column or single-factor analysis (Brown-Forsythe test), and two-way ANOVA for grouped analysis using Graphpad Prism version 10 with p-value < 0.0001, unless stated. Fluorescent intensity was calculated by subtracting the background using ImageJ (NIH). For in vivo experiments, a minimum of 6 larvae was utilized per condition per experiment.

## Data availability

The authors declare that all supporting data of this study are available in the article and its supplementary information files. Additional information can be requested from the corresponding author upon reasonable request.

## Author contributions

GP and DB conceptualized the study. DB acquired the funding. GP carried out the investigation. BP carried out EMSA, AFM imaging, and quantification. HD carried out the studies in a zebrafish model. GP carried out imaging and analyzed the data. SD provided the facilities and supervised the in vivo experiments. GP wrote the original manuscript, which was reviewed and edited by all the authors.

