## Supporting information for "Membrane sialylation orchestrates cellular gateways: A spatiotemporal analysis of cellular transport using DNA nanocages via membrane charge modulation"

| Primer | Nucleotide sequence |
|---|---|
| **M1** | 5'-ACATTCCTAAGTCTGAAACATTACAGCTTGCTACACGAGAAGAGCCGCCATAGTA-3' |
| **M2** | 5'-TATCACCAGGCAGTTGACAGTGTAGCAAGCTGTAATAGATGCGAGGGTCCAATAC-3' |
| **M3** | 5'-TCAACTGCCTGGTGATAAAACGACACTACGTGGGAATCTACTATGGCGGCTCTTC-3' |
| **M4 (Cy3/Cy5 at 5')** | 5'-TTCAGACTTAGGAATGTGCTTCCCACGTAGTGTCGTTTGTATTGGACCCTCGCAT-3' |

***Table S1:*** *Nucleotide sequence of oligonucleotide primers used for DNA TDN synthesis.*

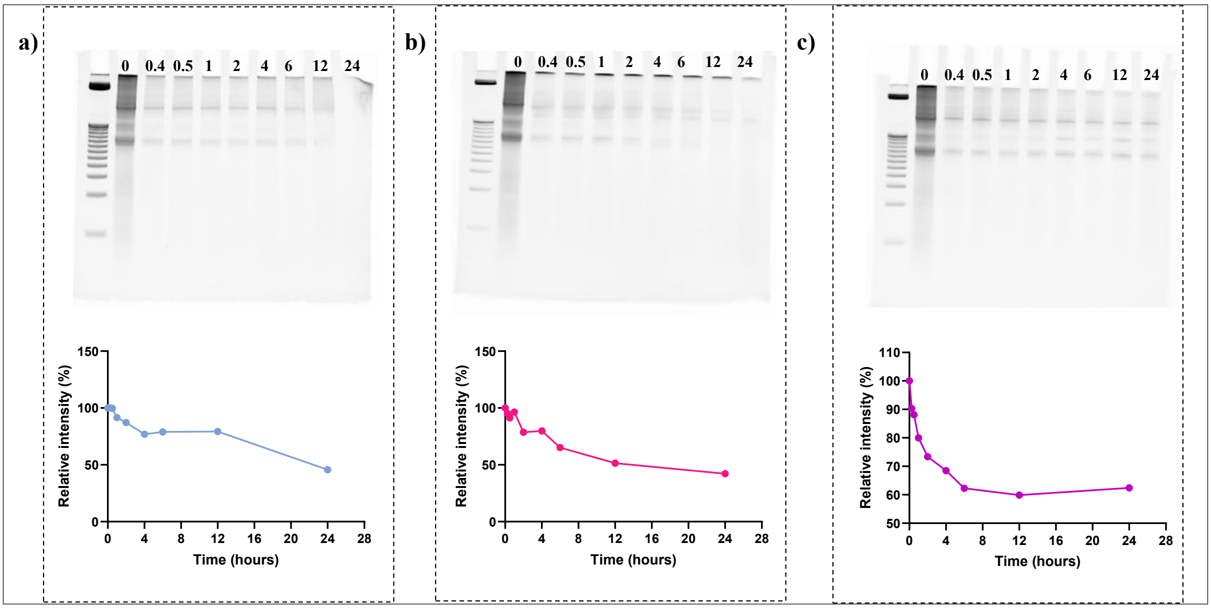

***Figure S1:*** *EMSA showing the stability of DNA TDN incubated in a) DMEM serum-free media (at 37ºC), b) 10% FBS (at 37ºC), and c) E3 media (at 28ºC) for 0.25, 0.5,1,2,4,6,12 and 24 hours. 10% PAGE was used, and band intensities were measured using ImageJ.*

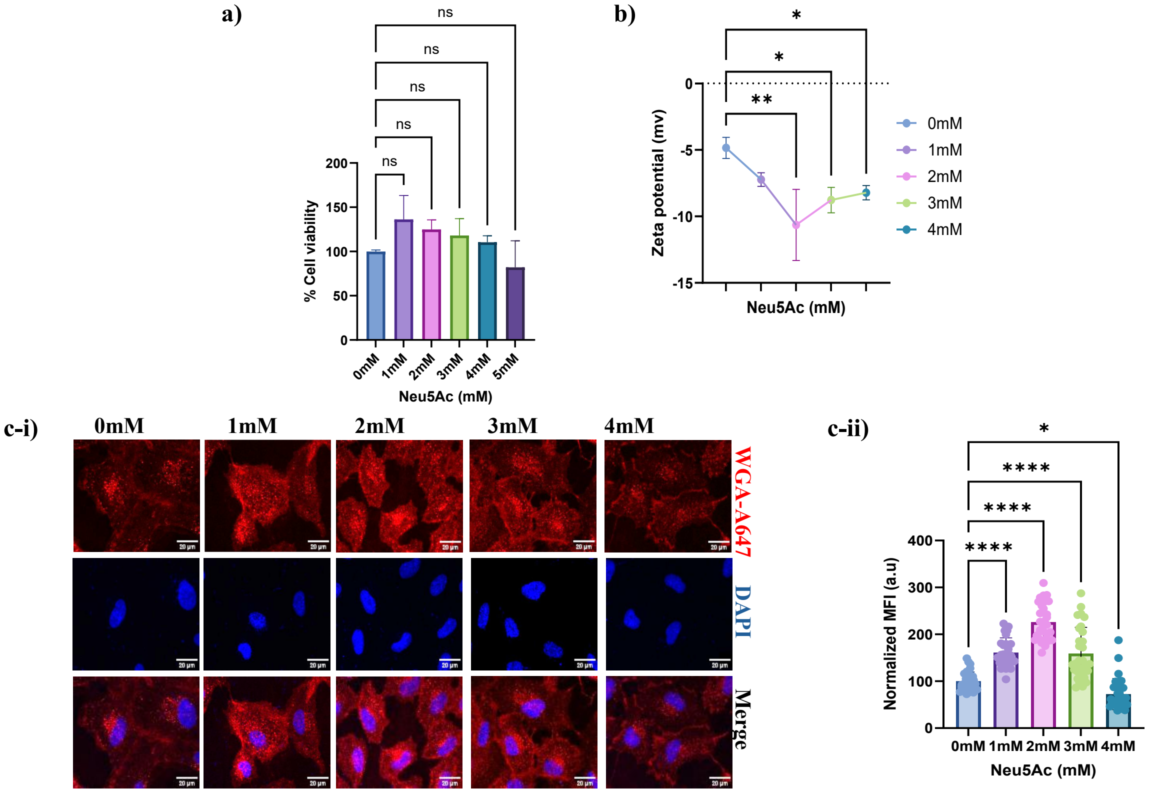


***Figure S2****: a) Cell viability assay (MTT assay) performed on RPE1 cells grown in DMEM complete media, supplemented with increasing concentrations of Neu5Ac (1mM to 4mM) for 96 hours at 37ºC and 5%CO*$_2$*. Cytotoxicity was not observed in the tested concentrations. b) Corresponding zeta potential measurement of RPE1 cells using Zetasizer Nano ZS analyzer, a higher negative charge was observed with 2mM Neu5Ac (Ordinary one-way ANOVA, p-value<0.005**), c) Confocal images of WGA staining on RPE1 cells grown in increasing concentrations of Neu5Ac (1mM-4mM). c-ii) Quantification of fluorescent intensity revealed increased WGA fluorescence at 2 mM Neu5Ac (p-value<0.0001****). Data plotted represent mean±s.d. N=30 cells were considered per treatment, N=3. Ordinary one-way ANOVA was used. Scale bar is 20µm.*

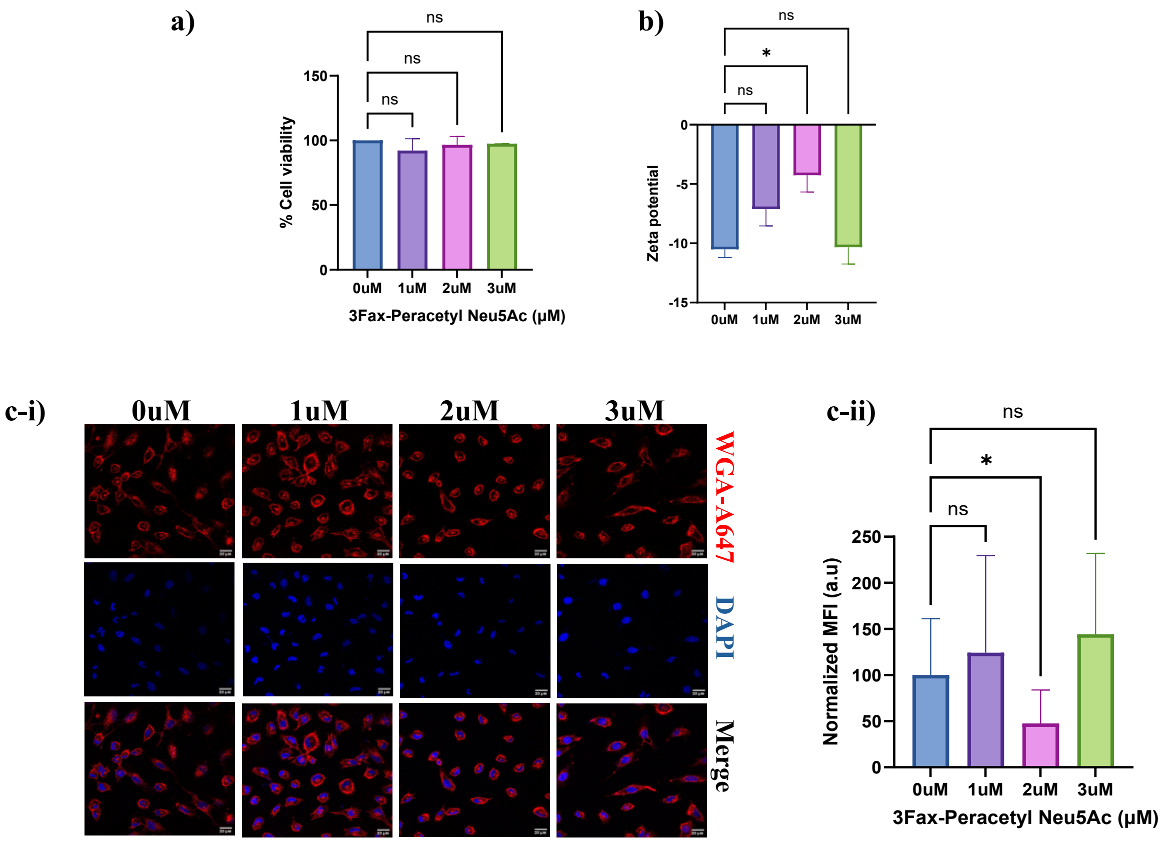


***Figure S3:*** *a) Cell viability assay performed on RPE1 cells grown in DMEM complete media, supplemented with increasing concentrations of 3Fax-Peracetyl Neu5Ac (1μM to 3μM) for 96 hours at 37ºC and 5%$CO_2$. Cytotoxicity was not observed in the tested concentrations. b) Corresponding zeta potential measurement of RPE1 cells using Zetasizer Nano ZS analyzer, a reduction in negative charge was observed with 2μM 3Fax-Peracetyl Neu5Ac (Ordinary one-way ANOVA, p-value<0.0215*), c) Confocal images of WGA staining on RPE1 cells grown in similar conditions. c-ii) Quantification of fluorescent intensity revealed a reduction in WGA fluorescence at 2μM 3Fax-Peracetyl Neu5Ac (p-value<0.0001****). Data plotted represent mean±s.d. N=30 cells were considered per treatment, N=3. Ordinary one-way ANOVA was used. Scale bar is 20μm.*

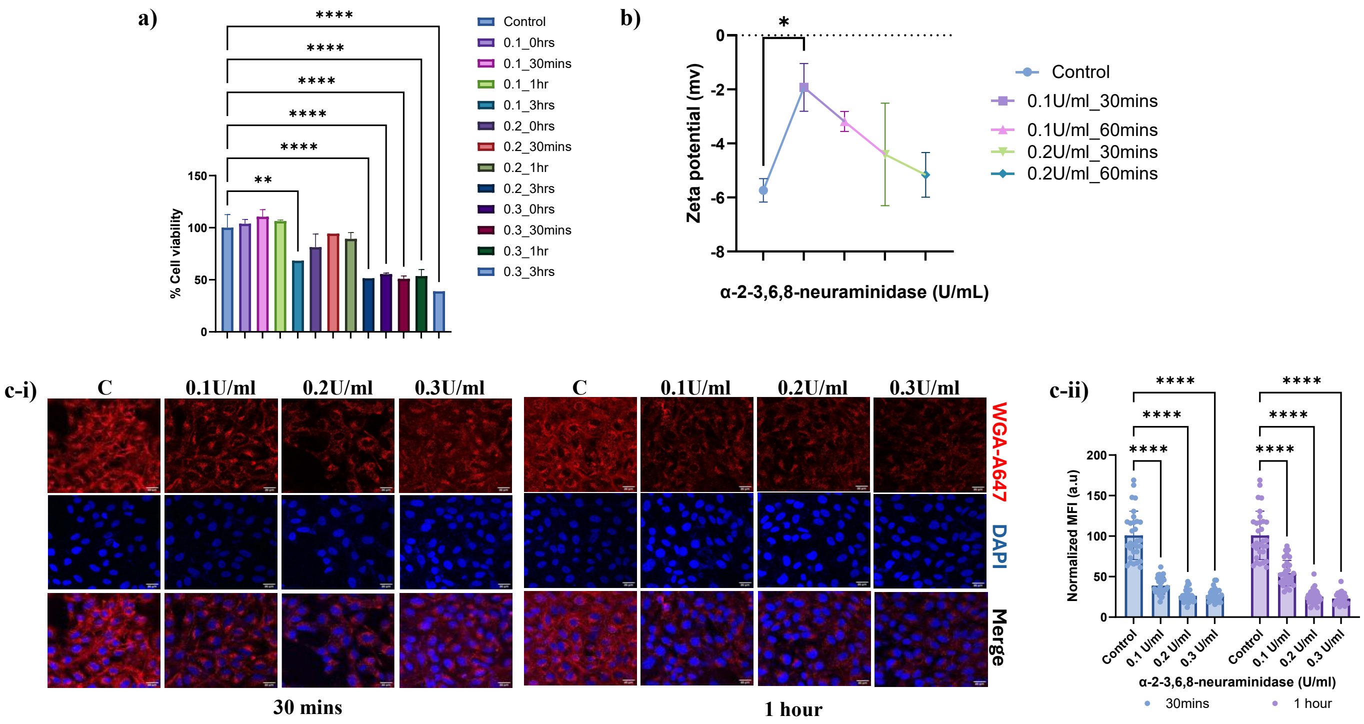


**Figure S4:** *a) Cell viability assay performed on RPE1 cells grown in DMEM complete media, supplemented with 2µM 3Fax-Peracetyl Neu5Ac for 96 hours at 37ºC and 5%$CO_2$, followed by treatment with α-2-3,6,8-neuraminidase (0.1,0.2 and 0.3 U/ml for 30minutes, 1 hour, and 3 hours). Cytotoxicity was observed at all tested concentrations after 3 hours of incubation, and at 0.3 U/ml enzyme concentration (Ordinary one-way ANOVA, p-value < 0.0001****). b) Corresponding to the zeta potential measurement of RPE1 cells using Zetasizer Nano ZS analyzer, a reduction in negative charge was observed with 0.1 U/ml enzyme treatment for 30 minutes (Ordinary one-way ANOVA, p-value<0.01*), c) Confocal images of WGA staining on RPE1 cells grown in similar conditions. c-ii) Quantification of fluorescent intensity revealed a reduction in WGA fluorescence at all tested enzyme concentrations and time (p-value < 0.0001****). Data plotted represent mean±s.d. n=30 cells were considered per treatment (N=3). Ordinary two-way ANOVA was used. Scale bar is 20µm.*

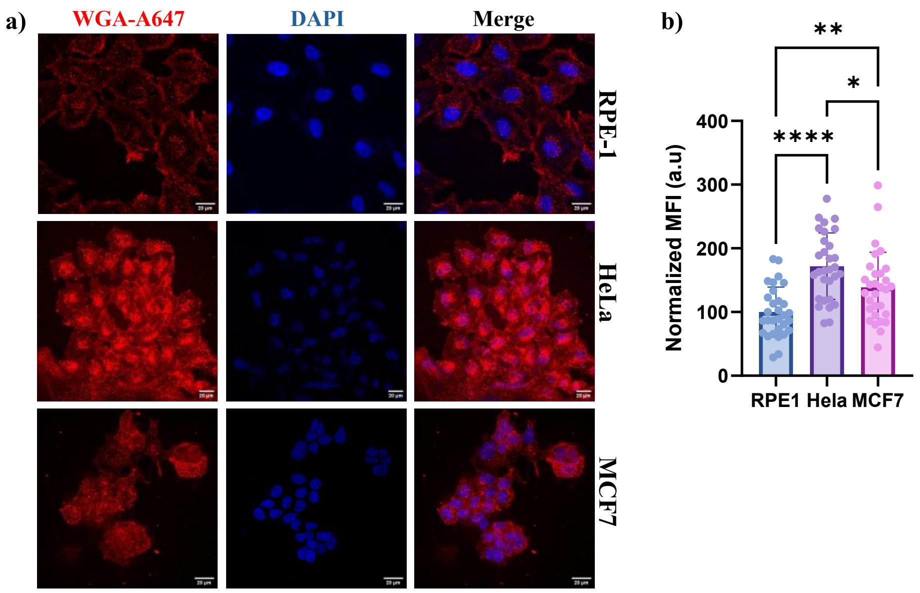


***Figure S5:*** *a) confocal images of WGA staining performed in RPE1, HeLa, and MCF7 cells. b) Quantification of fluorescence indicates an increase in WGA binding in cancerous cell lines (HeLa, MCF7) compared to the non-cancerous RPE1. Data plotted represent mean±s.d. n=30 cells were considered, N=3, One-way ANOVA (p-value <0.0001****).*

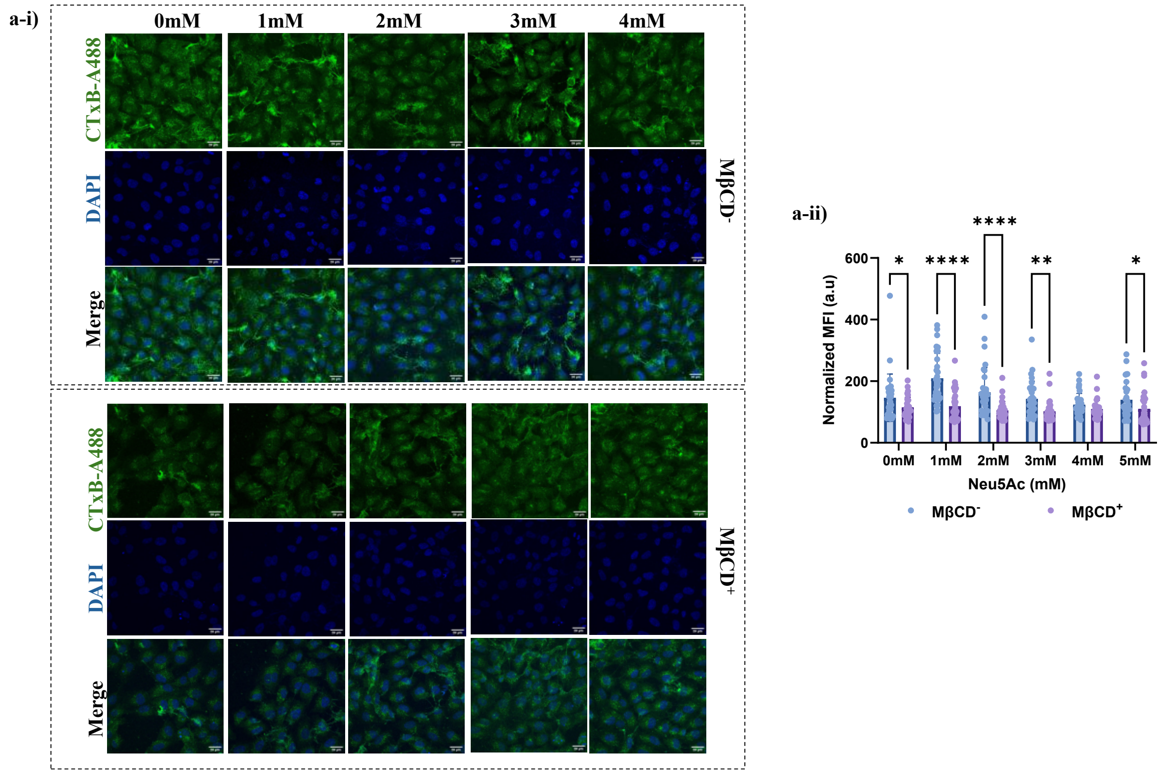


***Figure S6****: a) Confocal images of CTxB RPE1 cells grown in increasing concentrations of Neu5Ac (1mM to 4mM) in the presence and absence of MβCD. b) Quantification of fluorescent intensity revealed a significant reduction in CTxB fluorescence at the tested concentrations, with a higher reduction in 1mM and 2mM Neu5Ac. Data plotted represent mean±s.d. n=30 cells were considered, N=3, Two-way ANOVA (p-value <0.0001****).*

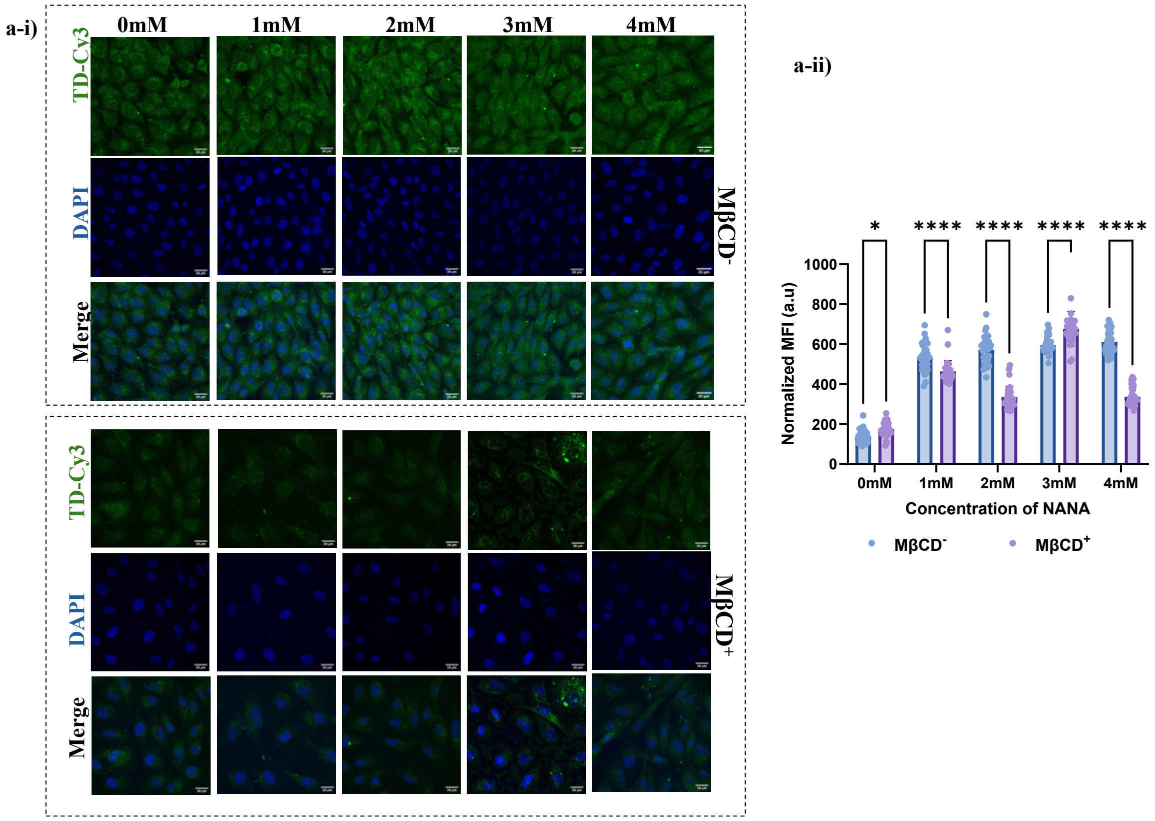


***Figure S7**: a) Confocal images of Cy3-TDN uptake in RPE1 cells grown with increasing concentrations of Neu5Ac (1mM to 4mM) in the presence and absence of MβCD. b) Quantification of fluorescent intensity revealed a significant reduction in TDN uptake at 1mM and 2mM Neu5Ac. Data plotted represent mean±s.d. n=30 cells were considered, N=3, Two-way ANOVA (p-value <0.0001****).*

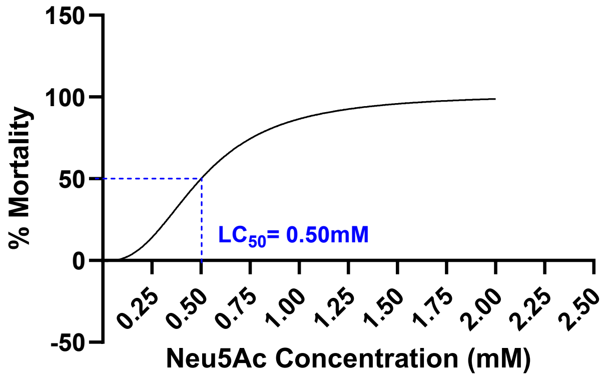


***Figure S8**: Cytotoxicity study of zebrafish larvae grown in increasing concentrations of Neu5Ac (0.25mM to 2mM) revealed an LC50 of 0.5mM.*

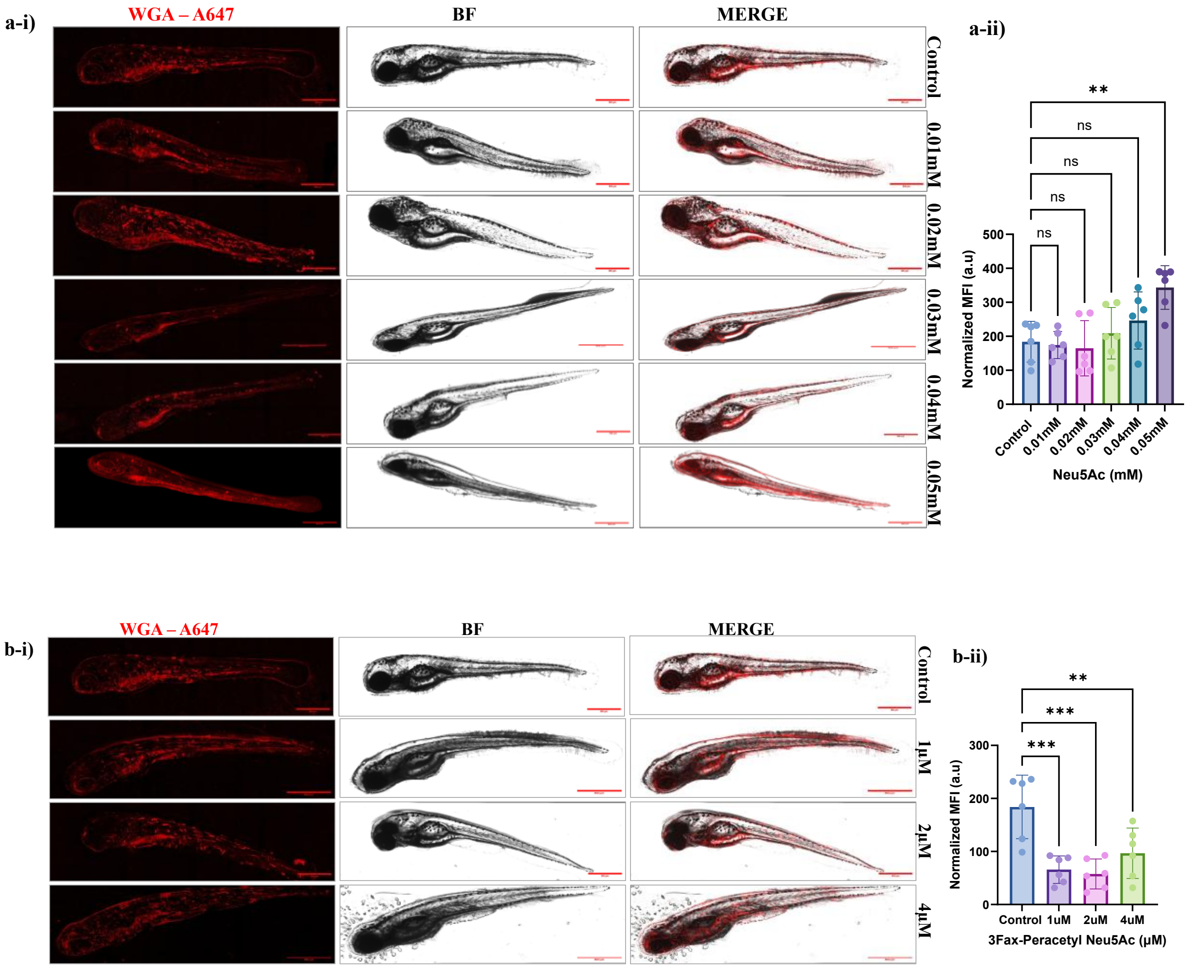


***Figure S9***: *Confocal images of WGA staining performed on zebrafish larvae at 96hpf grown in E3 media supplemented with a) increasing concentrations of Neu5AC (1mM to 4mM) and b) 3Fax-Peracetyl Neu5Ac (1 μM to 4 μM). Quantification of WGA fluorescence revealed an increase in binding at 0.05mM Neu5Ac (One-way ANOVA (p-value <0.0009****) and a significant reduction at 1 μM 3Fax-Peracetyl Neu5Ac. Data plotted represent mean±s.d. n=6 larvae were considered per condition, N=3, One-way ANOVA (p-value <0.0002****).*
